# DEMINERS enables clinical metagenomics and comparative transcriptomic analysis by increasing throughput and accuracy of nanopore direct RNA sequencing

**DOI:** 10.1101/2024.10.15.618384

**Authors:** Junwei Song, Li-an Lin, Chao Tang, Chuan Chen, Qingxin Yang, Dan Zhang, Yuancun Zhao, Han-cheng Wei, Kepan Linghu, Zijie Xu, Tingfeng Chen, Zhifeng He, Defu Liu, Yu Zhong, Weizhen Zhu, Wanqin Zeng, Li Chen, Guiqin Song, Mutian Chen, Juan Jiang, Juan Zhou, Jing Wang, Bojiang Chen, Binwu Ying, Yuan Wang, Jia Geng, Jing-wen Lin, Lu Chen

**Author notes:** Correspondence to (L. C.); (J-w. L.); (J. G.). These authors have contributed equally to this work (Junwei Song, Li-an Lin, Chao Tang, Chuan Chen).

## Abstract

Nanopore direct RNA sequencing (DRS) advances RNA biology but is limited by relatively low basecalling accuracy, low throughput, yet high RNA input and costs. Here we introduce a novel DRS toolkit, DEMINERS, which integrates an RNA multiplexing experimental workflow, a machine-learning barcode classifier based on Random Forest and a novel basecaller built on an optimized convolutional neural network providing an additional species-specific training module. With the increased accuracy in barcode classification and basecalling, DEMINERS can demultiplex up to 24 samples and the required RNA input and running time are both substantially reduced. We demonstrated the applications of DEMINERS in clinical metagenomics, cancer transcriptomics and parallel comparison of transcriptomic features in different biological conditions, revealing altered airway microbial diversity in COVID-19 and a potential role of m^6^A in increasing transcriptomic diversity in glioma and the mature blood-stage of malaria parasites. Overall, DEMINERS is a simple, robust, high-throughput DRS method for accurately estimating transcript levels, poly(A) lengths, and mutation and RNA modification heterogeneity at single-read level, with minimal sequencing biases.

## Introduction

The advance of third-generation sequencing technologies has revolutionized our ability to perform long-read sequencing for genomes and transcriptomes^1^. Oxford Nanopore Technologies’ (ONT) direct RNA sequencing (DRS) exemplifies this revolution, enabling the direct sequencing of DNA, RNA, and their modifications without fragmentation or amplification steps. This technology employs an array of protein nanopores in a synthetic membrane, allowing the sequencing of RNA molecules directly^2^. As motor enzymes ratchet a single RNA molecule through the pore, it causes electric current fluctuations when different nucleotides block the pore, a process that can be recorded to determine the RNA sequence using basecalling algorithms^3^.

DRS technology, capable of producing long sequencing reads up to 21 kb^4^, is powerful in obtaining full-length transcripts^5, 6^ and nearly complete RNA genomes^7, 8^. It is particularly effective in measuring RNA poly(A) tail length and analyzing the regulation function of these tails over gene expression^9–11^. Furthermore, DRS can detect RNA modification directly^12–15^, such as N6-methyladenosine (m^6^A), 7-methylguanosine (m7G)^16^, N1-methylpseudouridine^17^, pseudouridine (Ψ) and 2′-O-methylation (Nm)^15, 18^ and others, due to the unique current fluctuations these modifications produce.

The amplification-free library preparation method for DRS can sidestep amplification or RT bias^19^, making it particularly useful in identifying exitrons (exonic introns) that likely arise from reverse transcription artifacts^20^. DRS is particularly useful in identifying isoform-specific RNA structure^21^, tRNA^22, 23^ and rRNA^16^ modification, as well as in genome sequencing of RNA viruses^7, 8, 24, 25^. Another application of DRS is to analyze the dynamics of RNA metabolism by labeling nascent RNAs with base analogs (for example, 5-ethynyluridine^26^ and 4-thiouridine^27^). DRS has been applied in studies on (epi)transcriptome analyses across a wide range of species, including humans^4, 28^, animals (nematodes^6, 9^, insects^29, 30^etc), plants^31–33^, and microorganisms including bacteria^34, 35^, archaea^36^, yeast^37^, zooplankton^38^, viruses^11, 39–44^ and parasites^45–47^. Additionally, DRS has been applied in pathogen identification in clinical samples^24, 48, 49^.

Despite the advantages, to date DRS still faces several challenges. Firstly, DRS requires high RNA input which is especially challenging when dealing with rare or limited biological samples. Secondly, one RNA sample per flow cell leads to high cost and technical complexity, posing a substantial barrier in large-scale studies. On contrary, multiplexing samples in one flow cell will reduce the required RNA input, the effort in library construction and the sequencing cost without compromising data quality^50^. Moreover, the multiplexing strategy minimizes batch effects that arise from processing samples individually, leading to more consistent and reliable data across different sequencing runs. Current methods like Poreplex^51^ and DeePlexiCon^52^, limited to demultiplexing only four samples, highlight the need for improved methodologies to enhance throughput, reduce costs, and minimize technical variability in DRS applications.

However, basecalling during the demultiplexing process is quite challenging due to the lower sampling rate of RNA at 70 bases per second through nanopores, in contrast to 450 bases per second of DNA. This significant difference necessitates the development of specialized basecalling methods tailored to the distinct signal characteristics inherent to RNA. Currently, the median basecalling accuracy for DRS stands at about 86%^53^, necessitating a leap over the 90% threshold for high basecalling accuracy to reliably document genetic elements such as short exons and exon junctions^53^. This is especially crucial for RNA viruses and pathogens with high mutation rates. Although the utilization of CNNs has improved basecalling accuracy^54^, no studies have developed species-specific basecalling to mitigate the challenge of low accuracy in species-specific studies.

Here, we introduce DEMINERS (<u>D</u>emultiplexing and <u>E</u>valuation using <u>M</u>achine-learning Integrated in <u>N</u>anopor<u>e</u> direct <u>R</u>NA <u>S</u>equencing), an innovative machine-learning framework that significantly improved the efficiency and scalability for DRS (**Fig. 1**). DEMINERS employed a robust demultiplexing workflow that utilizes a Random Forest classifier to accurately demultiplex up to 24 samples per run. DEMINERS therefore reduces the required RNA input and sequencing costs, minimizing batch effects while maintaining high accuracy, thereby improving the consistency and reliability of comparative studies in RNA biology. Furthermore, DEMINERS incorporated a basecalling algorithm built on an optimized convolutional neural network (CNN) architecture, achieving state-of-the-art basecalling performance across different DRS datasets from animals, plants and microorganisms. Additionally, DEMINERS enables comprehensive downstream applications, including gene/isoform expression profiling, RNA variant and modification identification, assembly of RNA virus genomes and meta-genomic/transcriptomic analyses, providing reliable performance for diverse research purposes. Overall, DEMINERS improves the scalability and accuracy, while reduces the cost of DRS, facilitating the exploration of complex transcriptomic features in different organisms.

**Fig. 1.**
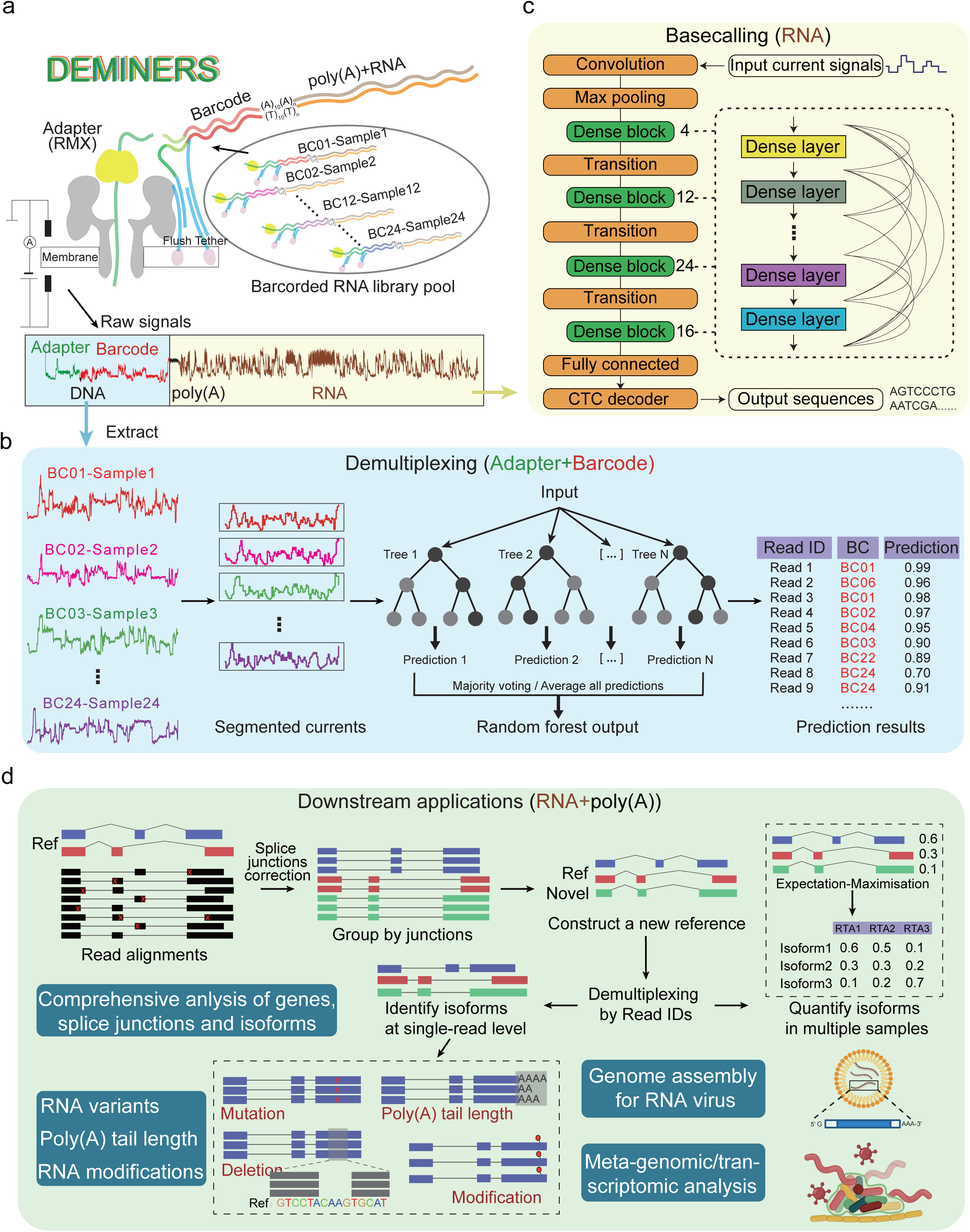
The overview of DEMINERS. **a,** Experimental workflow of barcoding and direct RNA sequencing pipeline. Each sample is ligated to an RNA transcription adapter (RTA) containing an RNA adaptor, a barcode (BC) and poly(T). These barcoded RNA samples are sequenced in a flowcell, producing raw signals of multiplexed barcodes and RNAs. **b,** Schematic illustration of DEMINERS machine-learning classifier based on Random Forest. The current signals of adapters and barcodes were extracted based on the distinct current changes introduced by poly(A) tails. The currents were then segmented into 100 segments/units according to the current changepoints. The normalized and segmented current signals were used as input for barcode classification based on random forest algorithm. **c,** Representation of DEMINERS basecaller built on an optimized convolutional neural network. The basecalling architecture employs an inter-layer connection strategy to foster feature reuse and mitigate the vanishing gradient. The basecaller incorporates a convolutional layer for denoising, followed by max pooling and 4 densely connected convolutional networks (Dense blocks) to decode raw current signals. The 4 dense blocks containing 6, 12, 24, and 16 dense layers, respectively. Then a fully connected layer with a log softmax activation is used for classification and a connectionist temporal classification (CTC) decoder outputs nucleotide sequence. **d,** Overview of downstream applications of DEMINERS. In this study, we show that the DRS reads retrieved by DEMINERS can be used for comprehensive analysis of genes, splice junctions, and isoforms. The splice junctions are corrected and grouped by junctions to construct a new reference. Isoforms are quantified using an expectation-maximization algorithm. By matching the demultiplexed read IDs, the mutations, deletions, poly(A) tail lengths and RNA modifications can be identified at the single-read level. Additionally, DEMINERS can perform genome assembly for RNA viruses and support meta-genomic/transcriptomic analysis.

## Results

### Overview of DEMINERS

To overcome the obstacles and improve the scalability of DRS, we developed DEMINERS, a comprehensive tool kit, that integrates three components: a multiplexing experimental workflow, a machine-learning classifier based on Random Forest and a novel basecaller built on an optimized CNN. The retrieved reads ensure comprehensive downstream analyses, including gene/isoform expression profiling, RNA variant and modification identification, genome assembly of RNA viruses, and meta-genomics/transcriptomics.

We first developed an experimental workflow for RNA multiplexing that integrates adapter-ligation and sample barcoding to ligate each RNA sample to a unique RNA transcription adapter (RTA) (**Fig. 1a**). The pooled RNA library is processed through a nanopore sequencer, generating raw electrical signals. For demultiplexing, DEMINERS utilized a Random Forest algorithm to classify the segmented current signals, allowing up to 24 samples to be demultiplexing simultaneously (**Fig. 1b**).

Next, to improve the accuracy of DRS basecalling, we reconstructed a convolutional basecaller inspired by DenseNet^55^, originally designed for image feature extraction. We adopted the reconstructed architecture to process one-dimensional electrical signals for basecalling, ensuring direct connection of each layer to the subsequent ones (**Fig. 1c**). Moreover, DEMINERS provides species-specific basecalling models to further increase the accuracy for diverse species (**Methods**).

For downstream analysis, we developed a comprehensive gene/isoform analysis workflow that facilitates the identification of novel transcripts by employing an expectation-maximization (EM) algorithm to estimate isoform abundance (**Methods**). DEMINERS also enables detection of RNA variations and modifications, genome assembly for RNA viruses and meta-genomic/transcriptomic analyses (**Fig. 1d**).

### Maximizing throughput and accuracy in demultiplexing

We designed and synthesized 48 RNA transcription adapters (RTAs), containing barcodes with lengths varying between 22–28 nucleotides (nt) (**Supplementary Data 1**). The design led to an increased Hamming distance amongst barcodes as their length extended, facilitating better differentiation (**Extended Data Fig. 1a**). These RTAs were then ligated to 51 in-vitro transcribed (IVT) RNAs and 5 DRS runs were performed, yielding a total of 6,276,168 valid reads uniquely aligned to the reference sequences. Considering the impact of sequencing depth on subsequent model training, we selected 24 barcodes with over 80,000 sequencing reads for further study (**Supplementary Data 1**). Notably, we observed that barcodes of mixed lengths significantly improved the performance of DEMINERS compared to using barcodes with uniformed 20-nt length (**Extended Data Fig. 1b**).

To advance the demultiplexing capability, we extracted signals of adapters and barcodes from the raw electrical signals based on the distinct current changes introduced by poly(A) tails. The optimal signal changepoints were identified according to the mean and variance of the normalized currents. The normalized currents were then segmented into small units, with the average current value of each segment/unit assigned as the feature values, which is used to train the classifier using different machine-learning algorithms (**Fig. 1c**, **Extended Data Fig. 1c**, **Methods**). This step retained the patterns of current signals while reduces the noises. We evaluated 6 different classification algorithms, including k-nearest neighbor (KNN), neural network (NNET), naïve Bayes (NB), classification and regression tree (CART), AdaBOOST and Random Forest (RF) to classify 4 barcodes in the test datasets (**Supplementary Data 1**). After plotting Receiver Operator Characteristic (ROC) and precision-recall (PR) curves for the 6 classifiers, we found that the area under the ROC curve (AUROC) were all above 0.93 (**Extended Data Fig. 1d**) and the area under the precision-recall curve (AUPRC) all greater than 0.83 (**Extended Data Fig. 1e**). Notably, RF outperformed the remaining methods in all parameters we measured (**Extended Data Fig. 1f**, **Supplementary Table 1**), achieving the highest AUROC of 0.9961, the highest AUPRC of 0.9906 and the highest accuracy of 95.66% (**Extended Data Fig. 1d-f**). The barcode classification of DEMINERS was thus built on RF algorithm.

Next, we evaluated the performance of DEMINERS in classifying different numbers of barcodes (from 2 to 24) and found the AUROCs were all higher than 0.99 (**Fig. 2a**). Moreover, AUPRCs were higher than 0.98 when classifying 10 barcodes, but still reached 0.95 when classifying 24 barcodes (**Fig. 2b**, **Supplementary Table 2**). We then assessed the accuracy and recovery rates of different numbers of barcodes (**Fig. 2c**). For instance, with a predicted probability cutoff set-off of 0.5, DEMINERS achieved an accuracy of 99.3% and an 89.2% recovery rate when classifying 10 barcodes, and 99.4% accuracy and 73.3% recovery for 24 barcodes (**Fig. 2c**, **Extended Data Fig. 2a**, **Supplementary Data 1**).

**Fig. 2.**
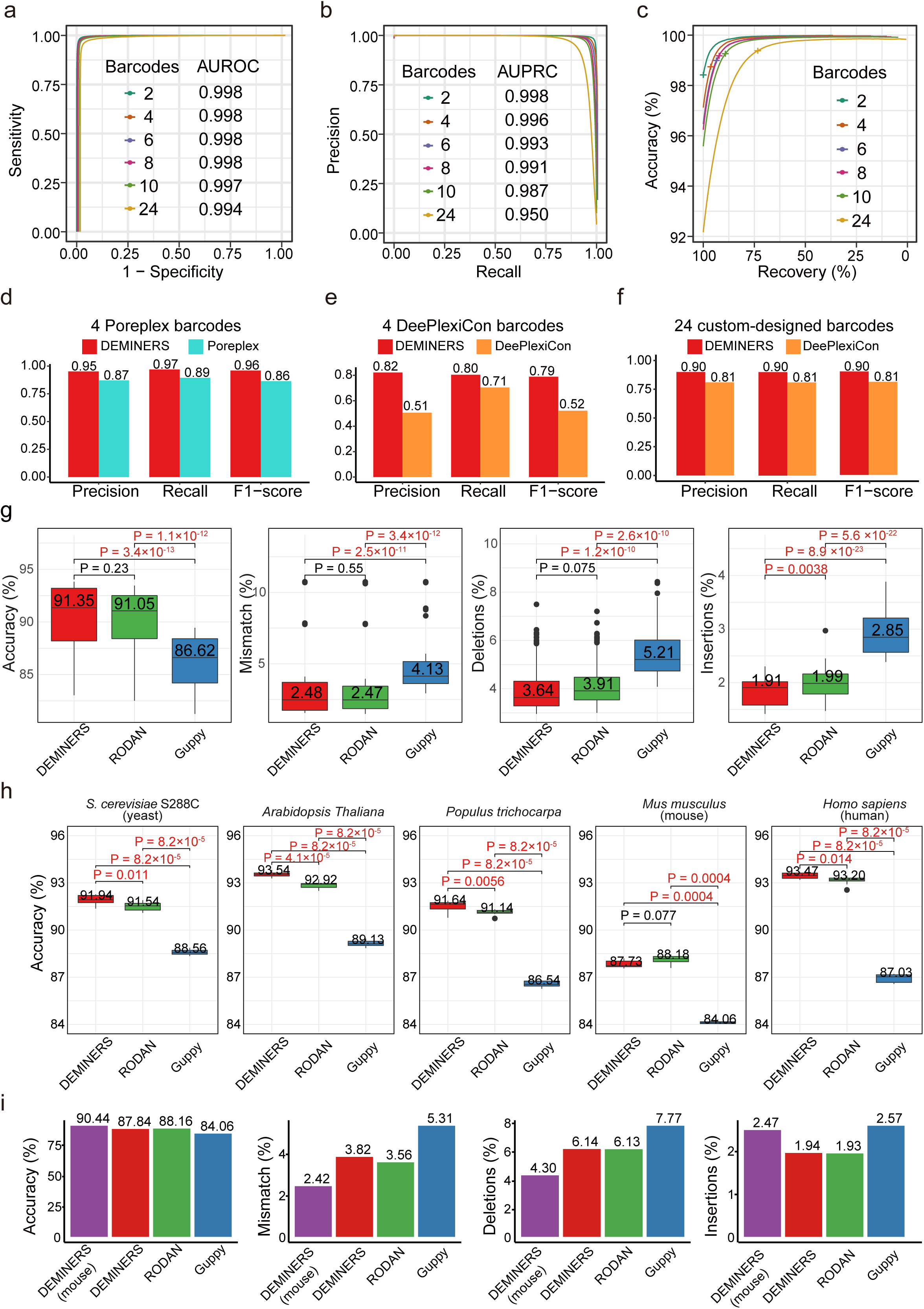
Performance of different demultiplexing and basecalling methods. **a-b,** Receiver Operating Characteristic (ROC) and Precision-Recall (PR) curves curves showing the performance of DEMINERS in demultiplexing direct RNA-seq (DRS) data generated with 2 to 24 barcodes. AUROC is the area under the ROC curve, and AUPRC is the area under the PR curve. **c,** Accuracy and recovery rates of DEMINERS in demultiplexing DRS data with different numbers of barcodes. The cross symbol represents the cutoff of predicted probability is at 0.5. **d,** Bar charts representing the precision, recall, and F1-score of DEMINERS and Poreplex^57^ classifying 4 Poreplex barcodes. **e,** Bar charts representing precision, recall, and F1-score of DEMINERS and DeePlexiCon classifying 4 DeePlexiCon barcodes. **f,** Comparison of DEMINERS and DeePlexiCon^58^ classifying 24 custom-designed barcodes. **g,** Box plots of the accuracy, mismatch, deletion, and insertion rates of DEMINERS, RODAN^64^, and Guppy in basecalling of the 10-species test set. The boxes show the median and lower/upper quantile, the dots indicate the outliers, and the P values were determined by Wilcoxon test. P values <0.05 are highlighted in red. **h,** Box plots showing the accuracy of DEMINERS, RODAN, and Guppy in basecalling DRS data of 5 different species. Each dataset was run for 8 times to ensure robustness, and P values were determined by Wilcoxon test. P values <0.05 are highlighted in red. **i,** Bar chats representing the accuracy, mismatch, deletion, and insertion rates of DEMINERS (mouse-specific and general modes), RODAN, and Guppy.

Furthermore, we conducted comparison analysis of DEMINERS with two existing methods, Poreplex^51^ and DeePlexiCon^52^. The evaluation was carried out using DRS datasets generated with the 4 barcodes originally designed for Poreplex or DeePlexiCon, and 24 barcodes designed in this study (**Supplementary Data 2**). When comparing DEMINERS to Poreplex using Poreplex 4-barcode dataset, the classification accuracies of DEMINERS ranged from 95.82 to 97.83% (mean ± standard deviation: 96.98±0.84), whereas the accuracies of Poreplex ranged from 83.4 to 93.18% (89.4±4.21) (**Extended Data Fig. 2b**). When demultiplexing DeePlexiCon 4-barcode dataset, the classification accuracies of DEMINERS were 90.57 to 94.43% (92.29±1.60), much higher than that of DeePlexiCon, ranging from 51.33 to 85.22% (68.39±15.16) (**Extended Data Fig. 2c**). Moreover, DEMINERS excelled DeePlexiCon and Poreplex in parallel comparisons in all measured characteristics, including precision, recall and F1-score (**Fig. 2d-f**), as well as accuracy, sensitivity and specificity (**Extended Data Fig. 2d-f**, **Supplementary Table 3**). When demultiplexing 24 barcodes, DEMINERS achieved accuracies between 83.24 and 93.78% (89.86±0.03), higher than the accuracies of DeePlexiCon, ranging from 69.67 to 87.97% (80.61±0.05) (**Extended Data Fig. 3a**). Although the sensitivity and specificity of DEMINERS and DeePlexiCon were comparable (**Extended Data Fig. 3b**), DEMINERS significantly outperformed DeePlexiCon in terms of precision, accuracy and recovery (**Extended Data Fig. 3c-d**). We further assessed the running time of DEMINERS and DeePlexiCon when processing the same amount (1-150 K) of DRS reads and found that the CPU running time of DEMINERS was around 1/12 of DeePlexiCon CPU time and 1/9 of DeePlexiCon GPU time (**Extended Data Fig. 3e**). In summary, DEMINERS effectively classified up to 24 barcodes with high accuracy and precision, outperforming existing methods like Poreplex and DeePlexiCon in accuracy, precision, recall and running time.

### Improving the basecalling accuracy

To improve the accuracy of DRS basecalling, we reconstructed a convolutional architecture inspired by DenseNet^55^ and provided a training module for species-specific basecalling (**Methods**). The basecaller of DEMINERS incorporated convolutional layers for denoising, max pooling for down-sampling, 4 dense blocks with transition layers for feature processing, and connectionist temporal classification (CTC)^56^ loss for gradient descent (**Fig. 1c**). In addition, we employed a memory optimization technique^57^ that significantly reduces memory consumption during model training, exemplified by reducing GPU memory usage from 6628 MB to 4700 MB with a batch size of 32 and a chunk size of 4096.

To evaluate the accuracy, we first trained the basecaller on a dataset containing 4 species (including human, *C. elegans*, *E. coli* and *Arabidopsis thaliana*) previously used by RODAN^54^. For the test set, we used an integrated 10 species dataset that includes 2 previous published datasets^11, 54^ and the in-house datasets, including four viruses [SARS-CoV-2, Porcine reproductive and respiratory syndrome virus (PRRSV), Seneca Valley virus (SVV), Porcine epidemic diarrhea virus (PEDV)], two plants (*A. Thaliana* and *Populus trichocarpa*), one fungus (*S. cerevisiae*), one eukaryotic parasite (*Plasmodiun berghei*) and two mammals (human and mouse) (**Supplementary Table 4**). The accuracy of basecalling the test sets of each species was evaluated by DEMINERS, RODAN^54^ and ONT-Guppy. We found that DEMINERS and RODAN significantly outperformed Guppy with higher accuracy, and lower mismatch, deletion and insertion rates (**Fig. 2g**). Compared to RODAN, DEMINERS had comparable accuracy, and mismatch and deletion rates, but fewer insertions (**Fig. 2g**).

Notably, DEMINERS showed higher basecalling accuracy than RODAN for 4 species, including *S. cerevisiae*, *A. Thaliana*, *P. trichocarpa* and human, except for mouse (**Fig. 2h**, **Supplementary Table 4**). To further improve the accuracy, we integrated a species-specific model training mode in DEMINERS. Using the RODAN mouse dataset to train the mouse-specific model, the basecalling accuracy for DEMINERS was improved from 87.84% to 90.44%, surpassing 88.16% accuracy of RODAN and 84.06% of Guppy (**Fig. 2i**). The mismatch and deletion rates were lower in DEMINERS mouse-specific model, but the insertion rate was higher (**Fig. 2i**, **Supplementary Table 4**). Moreover, although the mapping rates were comparable, the basecalled lengths were 11-bp longer using DEMINERS mouse-specific mode (517 bp) compared to RODAN (506 bp) (**Extended Data Fig. 3f**). Overall, DEMINERS demonstrated higher basecalling accuracy, longer read length and lower mismatch rates compared to Guppy and RODAN, especially after integrating a species-specific model, leading to improved performance in basecalling species-specific datasets and facilitating transcript assembly of the non-model organisms.

### Accurate pathogen identification, variant calling and genome assembly of RNA virus from multiplexed RNA samples

To assess whether DEMINERS can correctly distinguish different species and call variants from multiplexed RNA samples, we tested the performance of DEMINERS on three experimentally pooled RNA samples (**Fig. 3a**), containing 3 or 5 different species (**Supplementary Data 3**). The following 8 pathogens were used in the study: RNA viruses [Seneca Valley virus (SVV), Porcine epidemic diarrhea virus (PEDV), Porcine reproductive and respiratory syndrome virus (PRRSV), and Getah virus (GETV)], bacteria (*E. coli* and *S. enteritidis*), fungus (*S. cerevisiae*) and a parasite (*P. berghei*). For all three experiments, DEMINERS correctly identified the species with AUROC above 0.98 (**Fig. 3b**) and AUPRC above 95% (**Fig. 3c**), while achieving accuracy above 96% (**Fig. 3d**). Next, we merged the 3 datasets and successfully identified all species by metagenomic analysis (**Fig. 3e**) and found that the genomes of RNA viruses were significantly longer than transcripts of nonviral microorganisms (RNA viruses, mean: 2093; nonviral microorganisms, mean: 880; P < 2.2e-16, Wilcoxon test). These results demonstrated the reliability of our method and its potential in meta-genomic/transcriptomic applications.

**Fig. 3.**
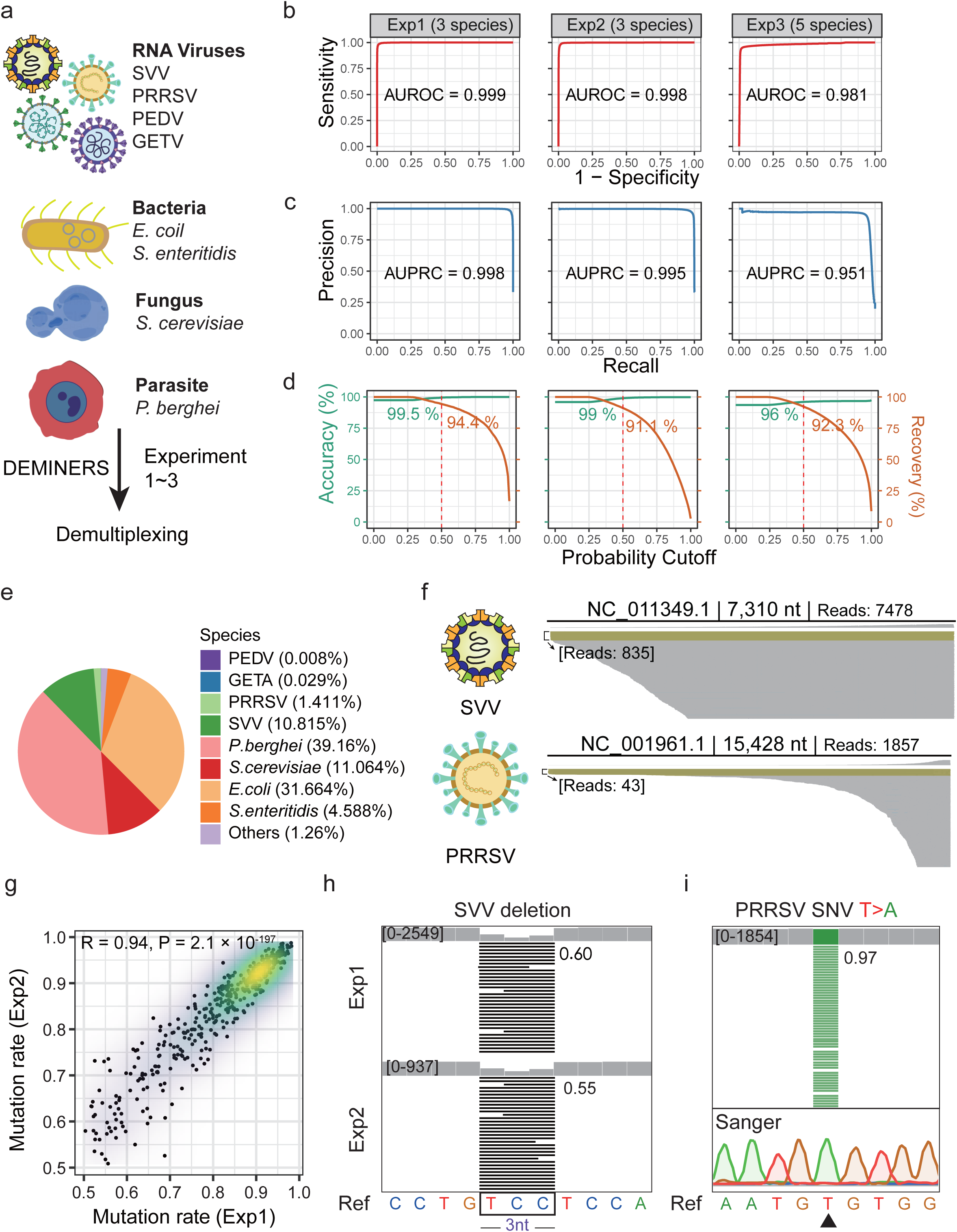
Performance of DEMINERS in pathogen identification, variant calling and genome assembly of RNA virus from multiplexed RNA samples. **a,** Experimental design to evaluate demultiplexing performance of DEMINERS. The multiplexed samples containing RNA isolated from various pathogens, including RNA viruses, bacteria, fungus, and parasite. **b-c,** Receiver Operating Characteristic (ROC) and Precision-Recall (PR) curves of DEMINERS in demultiplexing the samples of three DRS experiments (Exp). AUROC, the area under the ROC curve; AUPRC, the area under the PR curve. **d,** The accuracy and recovery rates at various predictive probability cutoffs. The red dashed lines indicate a predictive probability cutoff of 0.5. **e,** Pie chart depicting the distribution of species identified in pseudo-metagenomic analysis of the combined 3 DRS datasets. **f,** Integrative genomics viewer (IGV) visualizes genome coverage of Seneca Valley virus (SVV) (7,310 nt) and Porcine Reproductive and Respiratory Syndrome virus (PRRSV) (15,428 nt). Reads longer than 7000 nt for SVV and longer than 15,000 nt for PRRSV are colored in yellow and the number of reads were shown in brackets. **g,** Reproducibility assessment of single-nucleotide variants (SNVs) identified from demultiplexed SVV reads in two DRS experiments (Exp1 and Exp2). Pearson correlation coefficient (R) and relative P value were shown. **h,** IGV visualization of a 3-nt deletion (4022 to 4024) in SVV genome identified by DEMINERS in two experiments. The deleted sequences were boxed in the reference sequence (Ref). The numbers represent the average deletion frequencies. **i,** IGV visualization of a SNV (T-to-A at position 15307) in PRRSV genome identified by DEMINERS. The reference sequence (Ref) and Sanger sequencing chromatograms depicting the T-to-A variant (arrowhead) were shown. The number represents the frequency of the ‘A’ variant.

Next, we assessed whether DEMINERS retrieved-DRS sequences can be used for genome assembly of RNA virus. For SVV (genome 7,310-nt in length) and PRRSV (15,428-nt) where the sequence depths were adequate, the constructed genomes had 96.5% and 95.6% coverage with 96.1% and 95.6% identity compared to their reference genomes, respectively. Notably, there were 835 reads longer than 7,000-nt aligned to the SVV genome and 42 reads longer than 15,000-nt aligned to the PRRSV genome (**Fig. 3f**, **Extended Data Fig. 4a**), showing the capability of DEMINERS in assembly of RNA virus genomes.

Since RNA viruses have high mutation rates, we further analyzed the single-nucleotide variants (SNVs) in their genomes. For SVV, 414 and 423 SNVs were identified in two sequencing experiments (Exp1 and Exp2), of which 96.48% (411) were common with a high correlation rate (**Fig. 3g**, Person correlation R=0.94). For example, a C-to-V mutation and a 3-bp deletion of SVV were identified in both experiments (**Fig. 3h**, **Extended Data Fig. 4b**), indicating the high consistency between our multiplexed DRS experiments. Next, we aim to assess the accuracy of the identified SNVs. A total of 145 SNVs in PRRSV genome with high sequencing depth were all validated by Sanger sequencing (**Fig. 3i**, **Supplementary Data 3**). Combined, DEMINERS can accurately demultiplex experimentally pooled RNA samples, identify pathogens, call variants with high specificity and precision, and assemble genomes of RNA viruses.

### Unveiling altered respiratory tract microbial diversity in COVID-19

To test whether DEMINERS can be applied in clinical metagenomics, we collected 11 nasopharyngeal and 13 oropharyngeal swabs from 24 individuals infected with SARS-CoV-2. The total 24 samples were multiplexed and subjected to DRS following DEMINERS workflow, meanwhile NGS of SARS-CoV-2 genome enriched by multiplex PCR was performed (**Fig. 4a, Supplementary Data 4**). In total, 374,204 reads were retrieved after quality control, with an average length of 432-nt (ranging from 101- to 503,147-nt) (**Fig. 4a**, **Extended Data Fig. 4c-d**).

**Fig. 4.**
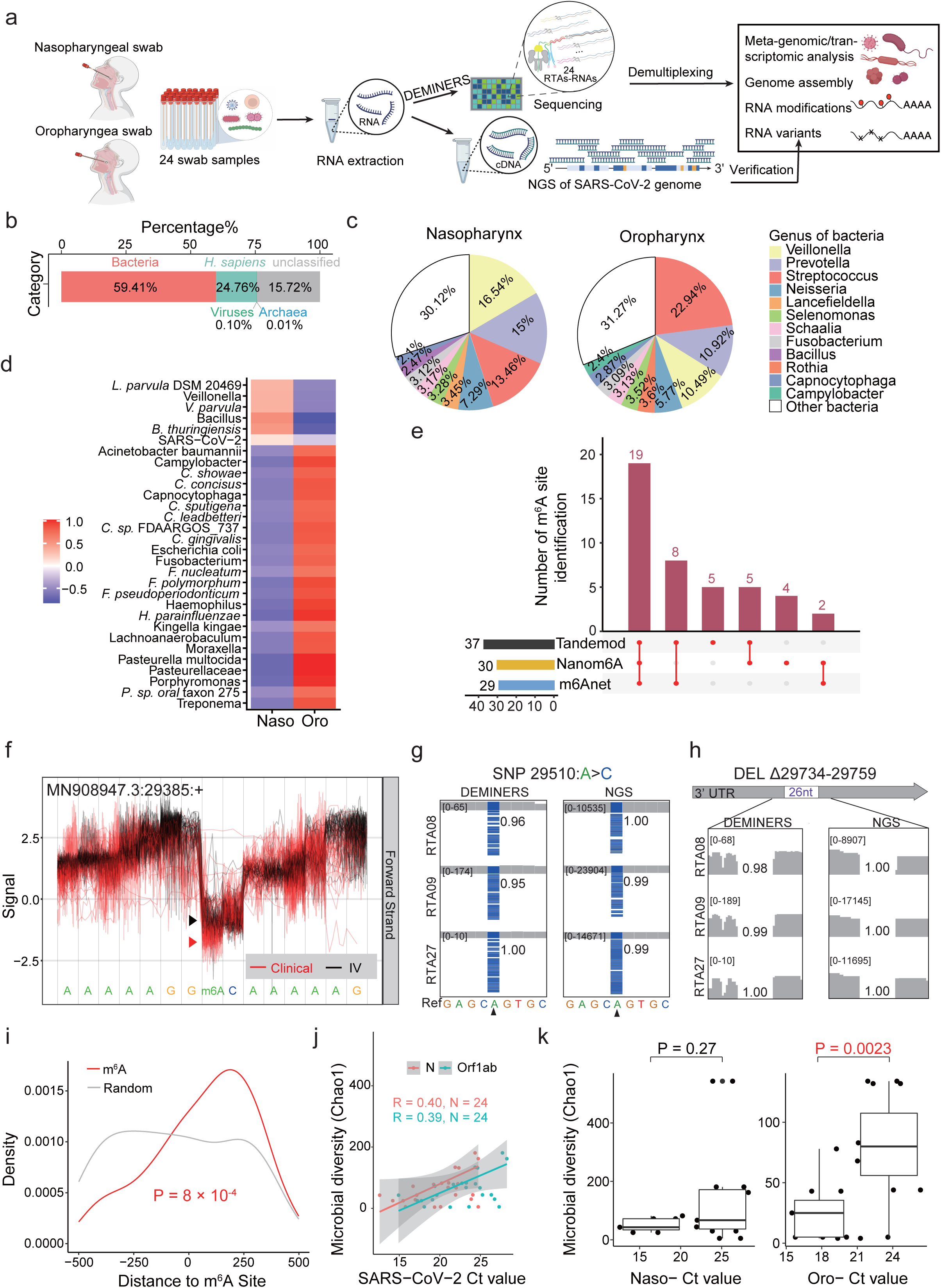
Metagenomic analysis of swab samples by DEMINERS. **a,** Study design. Eleven nasopharyngeal and thirteen oropharyngeal swabs were collected from 24 individuals infected with SARS-CoV-2. The isolated RNA were multiplexed and subjected to DEMINERS followed by metagenomics analysis. Meanwhile, the same individual samples were subjected to next generation sequencing (NGS) of SARS-CoV-2 genome enriched by multiplex PCR. **b,** Taxonomic classification analysis of the DEMINERS retrieved reads. **c,** Pie charts depicting the distribution of genus bacteria identified in the swab samples. **d,** Heatmap showing the differential distribution of microorganisms in nasopharyngeal (Naso) and oropharyngeal (Oro) swabs. **e,** UpSet plot showing m^6^A site overlaps among Nanom6A^14^, TandemMod^71^ and m6Anet^70^. The bar chart in lower left shows the total number of m^6^A sites identified by each software, and the lower right chart illustrates the counts of intersecting or unique sites identified by each software. **f,** Distinct ionic current signals indicating RNA modifications at position of 29,385 in SARS-CoV-2 genome. Red lines, SARS-CoV-2 from swab specimens (clinical); black lines, SARS-CoV-2 maintained in vitro culture (IV). **g,** IGV visualization of a SARS-CoV-2 SNV (A-to-C at position 29510, arrowhead) identified by DEMINERS and NGS in 3 swab samples. The numbers represent the read frequencies of the SNV. **h,** IGV visualization of a 26-nt deletion (29734 to 29759) in SARS-CoV-2 genome, identified by DEMINERS and NGS in 3 samples. The average deletion frequencies were shown. **i,** Density plot showing SNV densities flanking the identified m^6^A sites (red) and random sites (grey). P value, Wilcoxon test. **j,** Scatter plot showing Pearson correlation between Ct value of SARS-CoV-2 *N* and *Orf1ab* genes and microbial diversity (Chao1). R, Pearson correlation coefficient; N, relative sample size. **k,** Box plots showing the microbial diversity (Chao1) in nasopharyngeal (Naso-) or oropharyngeal (Oro-) swabs with high-Ct value (Ct>21) or low-Ct value (Ct≤21). P values were determined by Wilcoxon test.

Taxonomic classification analysis showed that around 60% reads belonged to bacteria, and the rest belonged to human (15.72%), viruses (0.1%) or archaea (0.01%) (**Fig. 4b**). A total of 377 genera were classified, 322 of which present in the nasopharynx and 329 in the oropharynx. While we observed a similar level of microbial diversity (measured by three methods, Chao1^58^, Shannon^59^ and Simpson^60^) (**Extended Data Fig. 4e-f**), the dominant bacterial genera were different (**Fig. 4c**). For instance, *Veillonella* (23,647 reads, 8.14%) was most prevalent in the nasopharynx, while *Streptococcus* was predominant in the oropharynx (8,326 reads, 11.09%) (P = 2.04e^−143^, Two proportion Z test) (**Fig. 4c, Supplementary Data 4**), consistent with a previous research on COVID-19 microbiome^61^. Principal component analysis (PCA) showed distinct clustering of the nasopharynx and oropharynx samples based on their microbial compositions (**Extended Data Fig. 4g**). Among the identified species, 30 of which showed differential distribution between the two sites (**Fig. 4d**). We also analyzed RNA modifications of the identified species. A total of 110 m^6^A sites were identified in *Streptococcus* using both m6Anet^62^ and TandemMod^63^ (**Extended Data Fig. 4h**) and these m^6^A sites were consistent with the DRACH motifs (**Extended Data Fig. 4i**).

Despite the low amount of viral RNA recovered from these clinical samples, we successfully mapped high-quality reads to the SARS-CoV-2 genome (mean quality 46.6) and assembled high-identity SARS-CoV-2 contigs (98.42%) (**Supplementary Data 4**). The reads mainly originated from the *N* and *ORF10* genes known to generate more subgenomic RNAs, consistent with a previous DRS study using SARS-CoV-2 maintained in Vero cell culture^11^ (**Extended Data Fig. 5a-b**). Using Nanom6A^13^, TandemMod^63^ and m6Anet^62^, we identified 43 m^6^A modification sites consistent with the DRACH motifs in the SARS-CoV-2 genome, 5 of which have been reported previously^11, 64^ (**Fig. 4e-f, Extended Data Fig. 5c-d**, **Supplementary Data 4**). Furthermore, we identified 26 point-mutations and two deletions (26- and 9-nt deletion), all validated by the parallel NGS (**Fig. 4g-h**, **Extended Data Fig. 5e**, **Supplementary Data 4**). An enrichment of SNVs around m^6^A peaks was found in our swab samples, consistent with the previous study on cultured SARS-CoV-2^11^ (**Fig. 4i**), indicating a potential relationship between mutation and m^6^A.

Notably, we found a positive correlation between the Ct values of SARS-CoV-2 PCR test and microbial diversity (R=0.4 and 0.39 for N and Orf1ab genes, respectively) (**Fig. 4j, Extended Data Fig. 5f**). Further analysis revealed that microbial diversity was significantly higher in oropharynx samples with high Ct-values (Ct>21) than those with low Ct-values (Ct≤21), but the diversity is comparable in the nasopharynx samples (**Fig. 4k**, **Extended Data Fig. 5g**).

Taken together, DEMINERS enabled DRS of clinical samples with low RNA amount and the reads retrieved can be used for metagenomics, variant calling and RNA modification analyses. Our analysis uncovered that SARS-CoV-2 abundance negatively impact on the oropharyngeal microbiota.

### Uncovering stage-specific transcriptomic features of the malaria parasite

The malaria parasite invades erythrocytes in the blood stage, classified into ring, trophozoite and schizont stages. The mature blood-stage parasites, schizonts withdraw from the circulating blood and cytoadhere to the microvasculature of internal organs which causes severe complications (**Fig. 5a**). To analyze the transcriptional and post-transcriptional features in different stages of malaria parasites, we performed DRS on 3 trophozoite RNA and 3 schizont RNA multiplexed and sequenced in one flow cell (**Supplementary Data 5**) and analyzed the expression level of genes and isoforms, poly(A) tail length of transcripts and m^6^A modifications (**Fig. 5a**).

**Fig. 5.**
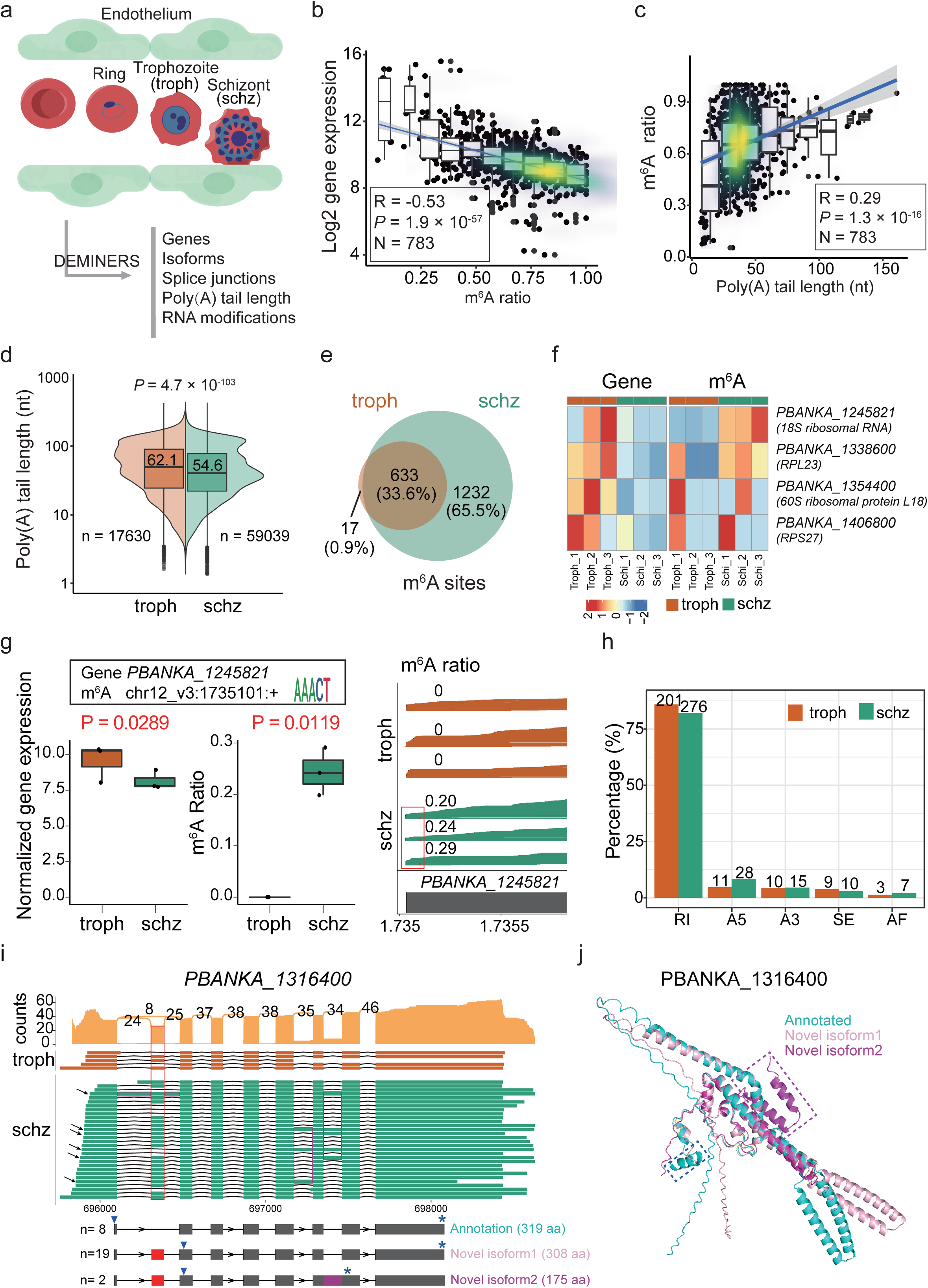
Parallel comparative analysis of transcriptomic features in different stages of malaria parasites. **a,** Schematics of the blood-stage malaria parasites. Trophozoite (troph) and Schizont (schz) stages of parasite were analyzed in this study using DEMINERS. **b-c,** Scatter plot showing Pearson correlation between m^6^A modification ratios and log2 gene expression or poly(A) tail length across all samples. R, Pearson correlation coefficient; P, P value; N, sample size. **d,** Violin plot showing the distribution of poly(A) tail lengths of all transcripts transcribed in trophozoite or schizont stages. The number of transcripts and the mean poly(A) tail length are shown. P value, Wilcoxon test. **e,** Venn diagram of m^6^A sites identified in trophozoites and schizonts. **f,** Heatmap illustrating the expression level of ribosome-related genes and their mean m^6^A modification levels in trophozoites and schizonts. **g,** Box plots showing the normalized gene expression of *PBANKA_1245821* and the ratios of m^6^A (chr12_v3:1735101) in trophozoites and schizonts. P values, Wald test for gene expression, T-test for m^6^A. Reads for gene, the ratio of m^6^A (marked in red) and PlasmoDB annotation are shown in the right. **h,** Bar charts showing the percentages and numbers of five major types alternative splicing events in trophozoites and schizonts. RI: retained intron, A5: alternative 5’ splice site, A3: alternative 3’ splice site, SE: skipped exon, AF: alternative first exon. **i,** Sashimi plot and read alignments for *PBANKA_1316400* in trophozoites and schizonts. The red boxes indicate the novel exon, the purple boxes indicate the retained introns, the arrows in the left indicate the reads of novel isoforms. Annotated transcript and novel isoforms with more than 2 DRS reads are shown below. The inversed triangles indicate the predicted start sites the stars indicate the predicted stop site and the numbers in the brackets showing the length of the predicted translated proteins. **j,** Pymol visualization of predicted protein structures of annotated or novel isoforms. The dashed blue box indicates the missing region of novel isoforms, and the dashed purple box highlights missing region of novel isoform 2.

Interestingly, we found that the gene expression level was significantly negatively correlated with m^6^A ratio (R= –0.53) (**Fig. 5b**), yet marginally correlated with poly(A) tail length (R= –0.15) (**Extended Data Fig. 6a**). In contrast, a positive correlation between m^6^A ratio and poly(A) tail length (R=0.29) was observed (**Fig. 5c**).

Notably, the poly(A) tail length and m^6^A sites differed between the two stages, with an average of 8-nt longer poly(A) tail in trophozoites (**Fig. 5d**), and 65.5% (1232) of the identified m^6^A sites present only in schizont, in sharp contrast to only 17 (0.9%) trophozoite-specific sites and 663 common sites (33.6%) (**Fig. 5e**, **Supplementary Data 5**). We further found that the genes upregulated in the developing stages, such as ribosome-related genes, were marked with m^6^A at the schizont stage, possibly leading to downregulation of these genes (**Fig. 5f**). For example, 18S ribosomal RNA *PBANKA_1245821* harbored a schizont-specific m^6^A site (chr12_v3 1735101) and its expression level was significantly downregulated at the schizont stage (**Fig. 5g**), indicating a negative regulation of m^6^A on gene expression in the mature stage.

Next, we investigated the isoforms expressed in trophozoites and schizonts. Intron retention (IR) was the most frequent alternative splicing type, accounting for 85.9% and 82.1% of all splicing events in trophozoites and schizonts, respectively (**Fig. 5h**). We identified 2,065 unannotated/novel isoforms in trophozoites and 2,510 in schizonts (**Extended Data Fig. 6b**), 22.5% (580) of the novel isoforms were stage-specific, mostly (502) schizont-specific (**Extended Data Fig. 6c**). Among the novel splice junctions (SJs), 18.2% and 51.2% were specific in trophozoites and schizonts, respectively (**Extended Data Fig. 6d-e**). Schizont stages showed both higher numbers of predicted novel isoforms and novel SJs, indicating that the isoform diversity is higher in this stage (**Extended Data Fig. 6b-e**). For example, for *PBANKA_1316400*, predicted to encode for a U1 snRNP-associated protein^65^, we identified an unannotated 80-bp exon, creating an alternative start site (**Fig. 5i**). The resulting new isoform is predicted to encode for a 11 amino acids shorter protein isoform, lacking an alpha helix at the N-terminus (**Fig. 5j**, **Supplementary Data 5**). Moreover, 4 introns of this gene showed IR events, generating 6 different types of isoforms, mainly in schizonts. One of schizont-specific IR isoform encodes for a small protein isoform lacking both N- and C-terminus (**Fig. 5 i-j**). For *PBANKA_0701800* (conserved protein with unknown function), we also identified a novel schizont-specific IR isoform (**Extended Data Fig. 6f**), introducing a premature stop codon possibly resulting in a non-coding RNA (**Extended Data Fig. 6g**, **Supplementary Data 5**). In addition, a novel exon-skipping isoform was identified for this gene, potentially encoding for a smaller protein isoform (**Extended Data Fig. 6g**).

Taken together, using DEMINERS, we analyzed stage-specific RNA biology of malaria parasites and identified schizont-specific features such as shorter poly(A) length, higher m^6^A modification levels and increased isoform diversity.

### Identifying m^6^A-associated isoforms in glioma

RNA modification and alternative splicing have been shown to play important roles in cancer progression and development^66–69^. To demonstrate the application of DEMINERS in analyzing RNA modifications in clinical cancer samples, we multiplexed 3 diffuse midline glioma-H3K27M mutant samples (DMG) and 1 glioblastoma sample (GBM) in one DRS run and classified the reads using DEMINERS (**Fig. 6a**). In addition, single-sample DRS (ss-DRS) was performed for 2 additional DMG and 3 GBM samples and NGS was also performed for verification (**Supplementary Data 6**).

**Fig. 6.**
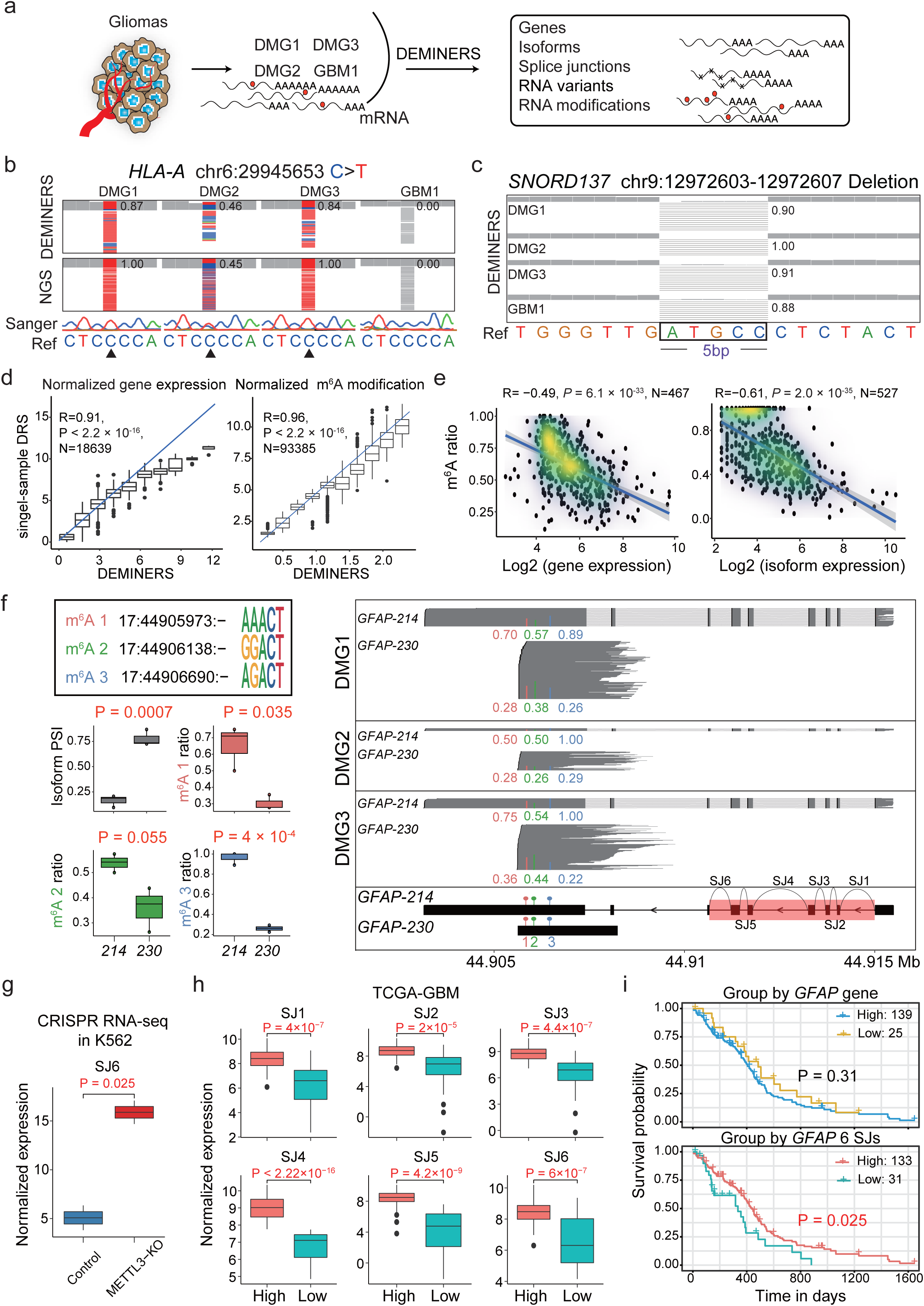
RNA variants, isoforms and m^6^A modification in human glioma. **a,** Schematic representation of multiplexed human glioma samples analyzed by DEMINERS. **b,** IGV visualization of a mutation (C-to-T at chr6 29945653 in *HLA-A*) identified by DEMINERS and NGS in all tumor samples. The numbers represent the read frequencies of the mutations. The Sanger sequencing chromatograms and the reference sequences (Ref) were shown. **c,** IGV visualization of a 5-bp deletion (12972603 to 12972607 at chr9 in *SNORD137*) identified by DEMINERS. The reference sequence and the average deletion frequencies are shown. **d,** Scatter plot showing Pearson correlation of normalized gene expression and m^6^A modification between DEMINERS and single-sample DRS. R, Pearson correlation coefficient; P, relative P value; N, sample size. **e,** Scatter plot showing Pearson correlation between m^6^A ratio and log2 gene or isoform expression. R, Pearson correlation coefficient; P, relative P value; N, sample size. **f,** Proportions of two isoforms (*GFAP-214* and *GFAP-230*) and their m^6^A ratios with motif at chr17 44905973, 44906138, and 44906690 in three DMG samples (**left**). P values, T-test. Total reads for *GFAP-214* and *GFAP-230* isoforms and m^6^A sites in DMG samples are shown (**right**). The colored dots represent the m^6^A sites and the numbers indicate ratios. The transcript structure based on Ensembl annotation is shown at the bottom, indicating the locations of the associated six splice junctions (SJs). The 6 SJs are located on chr17 with positions of 44914089−44915025 (SJ1), 44913824−44914027 (SJ2), 44913431−44913727 (SJ3), 44911798−44913268 (SJ4), 44911457−44911671 (SJ5), and 44910659−44911235 (SJ6). **g,** Box plot showing the normalized expression of SJ6 in METTL3-KO (n=2) and control (n=2) K562 cells (ENCODE CRISPR RNA-seq data^82^). P value, Wilcoxon test. **h,** Box plots showing normalized expression of 6 SJs of *GFAP-214* in high (n=133) and low (n=33) expression groups in the TCGA-GBM cohort^83^. P values, Wilcoxon test. **i,** Kaplan-Meier survival curves showing overall survival in the TCGA-GBM cohort. The survival curves were grouped by high (n = 139) and low (n = 25) expression of the *GFAP* gene (upper panel), or grouped by high (n = 133) and low (n = 31) expression of 6 SJs of *GFAP-214* (lower panel). P-values, log-rank test.

First, we examined whether DEMINERS can achieve high precision in identifying mutations and deletions. A total of 17,587 mutations were identified, and 97% of mutations (1,167) in GBM and 98% of mutations (11,791) of DMG were verified by ss-DRS (**Extended Data Fig. 7a**). For example, two DGM-specific mutations (chr6: 29945653:C>T and chr6: 29945609:C>G in *HLA-A*), one GBM-specific (chr16: 2519933:C>A in *ATP6V0C*) and one common mutation (chr17: 44906273:G>A in *GFAP*) identified by DEMINERS were all validated by NGS and Sanger sequencing (**Fig. 6b**, **Extended Data Fig. 7b-d**). Furthermore, the mutation rates were highly correlated between DEMINERS and NGS (R=0.93; **Extended Data Fig. 7e**, **Supplementary Data 6**). We also identified a 5-bp deletion mutation (chr9:12972603-12972607 in *SNORD137*) in all four glioma samples (**Fig. 6c**) using DEMINERS.

We compared the levels of gene expression and m^6^A modification identified by DEMINERS and ss-DRS. Although ss-DRS and DEMINERS were performed on different samples and lower read depth were obtained in the multiplexed samples, DEMINERS showed high correlation with ss-DRS in both gene expression (R=0.91) and m^6^A modification (R=0.96) (**Fig. 6d**, **Supplementary Data 6**). In detail, 99% of the 2,158 m^6^A sites were identified both by DEMINERS and ss-DRS, 39% of which were reported in m^6^A Altas^70^. The identified m^6^A modifications had DRACH motifs with an enrichment in the 3’UTR region (**Extended Data Fig. 7f**), suggesting that DEMINERS can correctly identify and quantify m^6^A level.

Next, we explored the relationship between m^6^A modification and the expression level of genes and isoforms. Interestingly, we observed that m^6^A ratio had a more prominent negative correlation to isoform expression level (R= –0.61) than the gene (R= –0.49) (**Fig. 6e**). For example, *GFAP* (glial fibrillary acidic protein), a marker of astrocyte differentiation and glioma diagnosis and prognosis^71^, had two isoforms, *GFAP-214* (a protein-coding isoform) and *GFAP-230* (with an unspliced intron, likely a ncRNA) detected in the three DMG samples. *GFAP-214* had a lower expression level compared to *GFAP-230* and exhibited significantly higher m^6^A modification at 3 sites (chr17 44905973, 44906138 and 44906690) (**Fig. 6f**). METTL3-mediated m^6^A modification have been reported to regulate cancer progression^72, 73^. We thus assessed whether the *GFAP-214* is regulated by the m^6^A writer, *METTL3*. A K562 *METTL3* knock-out (KO) dataset from ENCODE^74^ was used. We found that the splice junction 6 of *GFAP-214* was significantly upregulated in *METTL3*-KO cells (**Fig. 6g**), suggesting that the loss of METTL3 causes downregulation of m^6^A which leads to the upregulated expression of *GFAP-214*. To investigate the importance of the isoforms of *GFAP*, we analyzed the RNA-seq data of GBM in TCGA^75^. We found high expression of 6 SJs of *GFAP-214* associated with better prognosis (P= 0.025), whereas high- or low-expression levels of *GFAP* showed no significant difference in overall survival (P= 0.31) (**Fig. 6h-i**). Our findings showed that m^6^A potentially play a role in regulating the expression of *GFAP* isoforms, potentially contributing to cancer progression through complex post-transcriptional regulatory networks.

## Discussion

DRS enables direct sequencing of RNA molecules without fragmentation, reverse transcription or amplification, making it effective in obtaining information of full-length transcripts, poly(A) tail length, and RNA modifications. Despite its advantages, DRS faces challenges such as low throughput, low basecalling accuracy yet high input requirements. DEMINERS significantly improved the accuracies of barcode classification and basecalling, thereby increases the throughput of DRS and facilitates the complex analysis of RNA biology. Importantly, DEMINERS allows parallel comparisons of transcriptomic features in different biological conditions, increasing statistical power and reducing batch effects and sequencing cost.

Hitherto, the barcoding strategy offered by ONT is only suitable for DNA and cDNA sequencing, but not for DRS; and DeePlexiCon^52^ and Poreplex^51^ provide demultiplexing of only 4 barcodes. In this study we scaled up the demultiplexing capability to 24 samples while maintaining higher accuracy. We considered the GC content, melting temperature and secondary structure in the barcode design^76^. To ensure the accuracy of barcode classification, we opted for barcodes with different length and implemented a Hamming distance error correction algorithm^77^.

Next, we developed a novel signal smoothing method for the random forest algorithm, which preserved the major changes in the current signals while reduced the random noises by segmenting the current signals and speeding up the classification step. As a result, DEMINERS exhibits superior running speed, taking only 1/10 of the CPU time of DeePlexiCon^52^. Meanwhile, DEMINERS shows high flexibility, allowing preference for high classification accuracy or high read recovery by adjusting the classifier prediction probability.

The differences in current signals and translocation speeds between DNA and RNA, lead to a high error rate in basecalling. We developed a DRS-specific basecalling method reconstructed from DenseNet^55^, leveraging its advantages of facilitating information flow, feature reuse, and efficient gradient propagation. Meanwhile, we employed the memory optimization technique^57^, which significantly reduces memory consumption during model training. Moreover, we provide species-specific basecalling modes, further improving the accuracy of basecalling.

With improved barcode classification and basecallling, we demonstrated the application of DEMINERS in real-world samples, showcased its capability of metagenomics, genome assembly of RNA viruses, and analyses of transcripts, mutation, and RNA modification. The real-time ONT sequencing is particularly useful in rapid detection of infection, antibiotic/antimicrobial resistance and metagenomics^1, 78, 79^. Although compared to cDNA/DNA sequencing DRS provides additional information, such as RNA modifications. The technical difficulties remain for application of DRS in infection detection because of the low amount of viral RNAs in swab samples. It is estimated that only an average of 58 SARS-CoV-2 RNA copies/µL in nasopharyngeal swabs and 14 copies/µL in oropharyngeal swabs^80^. Thanks to the increased accuracy in barcode classification and basecalling, DEMINERS can demultiplex up to 24 samples and the required RNA input is thus substantially reduced. We demonstrated the application of DEMINERS in metagenomic analysis of 24 multiplexed nasal/ oropharyngeal swabs and revealed that the microbiota was altered in COVID-19, in line with previous studies^61, 81, 82^.

We demonstrated the application of DEMINERS in parallel comparison studies of different biological conditions without library construction nor sequencing batch effect. We not only identified many unannotated exons and spicing junctions, but also revealed stage-specific characteristics, including shorter poly(A) length, and higher transcript diversity and m^6^A modification in the mature stage of malaria parasites.

RNA modifications, particularly m^6^A methylation, have emerged as pivotal regulatory elements in microorganisms^83^, including viruses^84^, and cancers^85, 86^. Using DEMINERS, we not only accurately identified m^6^A modification sites, identified a negative correlation between m^6^A and gene expression in malaria parasites and gliomas alike to previous studies^87^, but also uncovered an even more prominent negative correlation between m^6^A and isoforms in gliomas. For example, m^6^A likely involved in splicing regulation of *GFAP* and is associated with glioma prognosis.

Together, DEMINERS brings together high accuracy barcode classifier based on machine-learning and improved baseballer based on a convolutional neural network, provides users with an economical solution for DRS, reducing both the RNA input and the sequencing cost and enables exploration of RNA biology in diverse biological processes.

## Online methods

### Generation of multiplexed direct RNA sequencing data

#### RNA transcription adapter design

A set of 40 RNA transcription adapters (RTAs) were designed, each comprising an RNA adaptor (RMX), a barcode, and a poly(T) sequence. In addition, the standard ONT RTA, 3 RTAs for DeePlexiCon^52^, and 4 RTAs for Poreplex^51^ were also included, making up a total of 48 RTAs were used in our study (**Supplementary Data 1**).

The barcode length ranged from 20 to 28 nucleotides (nt) increasing by 2-nt. We did not use ultra-long barcodes to prevent potential interference during the magnetic bead-based purification step removing free RTA. The OligoAnalyzer™ Tool was used to evaluate the GC content, melting temperature and secondary structure of DNA oligonucleotides to assess the quality of the DNA barcodes, The barcode sequences were listed in **Supplementary Data 1.**

#### Preparation of RTAs and in-vitro transcribed RNAs

Forward (1.54 μM) and reverse (1.4 μM) strands of the RTA sequences were mixed in the buffer (10 mM Tris-HCl pH 7.5, 50 mM KCl). The extra forward strands were used to avoid left-over reverse strands, which may reduce the yield of RNA-RTA products. The custom RTAs were annealed following the process: 95°C for 5 min, 65°C, 50°C, 37°C, and 22°C for 30 min at each temperature, then stored at 4°C. The formation of RT adapters was verified by 4% agarose gel electrophoresis.

Fifty-one in-vitro transcribed (IVT) RNAs, with lengths ranging from 177 to 748 bp, were generated for barcode testing and model training. The genes and primers used to generate cDNA are listed in **Supplementary Data 1**. The PCR products were determined by agarose gel electrophoresis. IVT RNAs were then generated by T7-RNA polymerase (NEB, M0251) using 1 μg of purified PCR products and purified using RNA Clean & Concentrator-5 kits (Zymo, R1015) following the manufacturer’s recommendations. The integrity and concentration of IVT RNAs were measured by Qsep100 (BiOptic Inc) and Nanodrop 2000 spectrophotometer (Thermo Fisher Scientific), respectively.

#### RNA multiplexing, library preparation and sequencing

RNA samples were ligated to pre-annealed custom RT adaptors (RTA) using NEBNext Quick Ligation Module (NEB, E6056) in separated reactions and reverse-transcribed by SuperScript IV Reverse Transcriptase (ThermoFisher, 18090200) to form RNA/DNA duplexes avoiding the formation of the secondary structure of RNA. The products were purified using 1.8X Agencourt RNAClean XP beads (Beckman, A63987). Next, equimolar amounts of reverse-transcribed RNA/DNA hybrids from each reaction were pooled and the RNA adaptors mix (RMX) were ligated to the 3′ end of RNA/DNA hybrid at room temperature for 30 mins. After ligation, the mixture was purified using 0.4X Agencourt RNAClean XP beads (Beckman, A63987), washed twice with the wash buffer, and eluted in the elution buffer. The RNA sequencing library was prepared using Direct RNA Sequencing Kit (SQK-RNA002, Oxford Nanopore Technologies) following the Direct RNA sequencing protocol (DRCE_9079_v2_revI_14Aug2019, ONT). The sequencing libraries were eluted in the RNA running buffer, loaded onto a primed R9.4.5 flowcell (ONT, FLO-MIN112 or FLO-PRO002), and sequenced on a MinION or PromethION sequencer (ONT) until all pores were inactivated. Five sequencing runs were performed with various combinations of RTA and IVTs and the details of read counts and quality were listed in **Supplementary Data 1**.

### Model training for demultiplexing DRS data

#### Basecalling and alignment

The raw current signal data from MinKNOW (Oxford Nanopore Technologies, v21.06.0) were used to obtain the RNA sequence data using Guppy (ONT, v6.5.7) with high-accuracy basecalling model (config: rna_r9.4.1_70bps_hac.cfg). Basecalled reads (FASTQ) were aligned to the reference sequences using minimap2^88^ (v2.15, -ax map-ont) with default parameters. The reference FASTA sequences were extracted from the reference genomes and listed in **Supplementary Data 1**. We filtered out the multiple alignments and the reads with mapping quality less than 60 for downstream analysis. The remaining high-quality and uniquely mapped reads were grouped based on the coupled RNA sequence, distinguished by mapped reference sequences.

#### Signal transformation

The raw current signals were extracted from the FAST5 files using R package rhdf5 (v2.30.1, https://github.com/grimbough/rhdf5). First, we implemented the standard double-exponential moving average (DEMA) algorithm from the R package smoother (v1.1, https://CRAN.R-project.org/package=smoother) to reduce the noise of raw current signals. In this step, we set the averaging period to 20 units, meaning that the average was calculated over 20 consecutive data points. Secondly, we extracted the signals of the adapter and barcode of each read from the denoised electrical signals, according to the characteristic higher and more stable current change generated by poly(A) tails and the lower current signals of DNA molecules. Specifically, we used the cpt.meanvar function of the R package changepoint^89^ (v2.2.2) to calculate the optimal change position from the first 20,000 denoised electrical signals which contains the adapter, barcode, and poly(A) signals for almost all reads. A single changepoint is denoted as the first observation of the new segment with a significant difference in the mean and variance using the cpt.meanvar function of AMOC^90^. Signals before the changepoint were extracted as adapters and barcodes, and at least 90% of adapter and barcode signals were smaller than the mean of poly(A) signals originating from the first stable segment after the changepoint. Finally, for the extracted adapter and barcode signals, the approximate (BinSeg) method^91^ of cpt.meanvar function was used to identify multiple changepoints with penalty. For each read, we identified 99 changepoints to divide the current signals into 100 units/segments and calculated the average current value of each unit as feature values.

#### Model training for demultiplexing

The feature values, defined from the mean current of 100 units, were used as predictors of each read. A matrix of 100 features generated from all reads was eventually used for model training, testing, and independent validation. The true label of each barcode signal was defined according to the RNA molecular which was pre-designed and linked to the barcode. To compare and identify optimal machine learning classification algorithms, we selected 4 barcodes from sequencing Run 4 as a test(**Supplementary Data 1**). For each barcode, we randomly selected 10,000 reads as a training set and the remaining reads were used as a test set. Model training was performed using the R package caret^92^ (v6.0-88). Six machine-learning algorithms were tested for barcode classification, including random forest (RF), naive Bayes (NB), K nearest neighbours (KNNs), bagged classification trees (CART), adaptive boosting classification trees (AdaBoost), and neural networks (NNet).

As RF algorithm outperformed other methods, it was selected for subsequent method development. We selected the 2, 4, 6, 8, 10 and 24 barcodes with the most reads from sequencing Run4 for RF training (**Supplementary Data 2**). Because the read numbers of each barcode differed, we randomly sampled 30,000 reads of each barcode from Run4 as a training set using the downSample function of R package caret^92^ (v6.0-88). The rest of the reads of the 24 barcodes were used as a test set and the reads from the Run3 were used as an independent validation set (**Supplementary Data 2**). To prevent overfitting, we applied the repeated K-fold cross-validation (10-fold, repeated 10 times) method to resample in the model training. To increase the computational efficiency, a parallel processing framework was employed in the model training using the R package doParallel (v1.0.16, https://CRAN.R-project.org/package=doParalle).

#### Method comparison

In the comparative study of DEMINERS and DeePlexiCon^52^ (v1.2.0), we randomly selected 24,000 reads (10,000 each from 24 barcodes with the most reads from Run4) to train the prediction models using DEMINERS and DeePlexiCon. Similarly, the rest of the reads of the 24 barcodes were used as a test set and the reads from Run3 were used as an independent validation set. In the comparison of DEMINERS and Poreplex^51^ (v0.5), because we did not have access to the raw data of Poreplex, and Poreplex (v0.5) does not support user-supplied data to train new models, we evaluated their pre-trained models on our data (**Supplementary Data 2**). We selected all reads of the 4 barcodes used by Poreplex (v0.5) from all sequencing runs, and randomly extracted 100,000 reads of each barcode as a training set. The remaining reads were used as a test set to evaluate and compare the performance of DEMINERS and Poreplex. For Poreplex, the classification of barcodes were performed using the recommended parameters^51^.

#### Performance Evaluation

The receiving operator characteristic curve (ROC) and the associated precision-recall curves were charted using the R package multiROC (v1.1.1, https://CRAN.R-project.org/package=multiROC), and the area under the curves (AUC) were calculated as AUROC and AUPRC, respectively. The relative accuracy, sensitivity, specificity, precision, recall and F1 score of each model were determined using the confusionMatrix function of R package caret^92^ (v6.0-88). The CPU time was calculated as the sum of the user time and system time evaluated using GNU time (https://www.gnu.org/software/time/).

### Basecalling model training

#### Architecture of the basecaller

In our architecture, three convolutional layers were used to denoise the signals, followed by a max pooling layer with a pooling size of 10 for downsampling. DEMINERS includes four dense blocks with 6, 12, 24, and 16 layers, respectively. Each dense layer starts with DenseNet typically started with a bottleneck layer that includes Batch Normalization (BN), a Sigmoid Linear Unit (SiLU)^93^, and then a 1×1 convolution to reduce dimensionality, followed by another BN, SiLU and a 3×3 convolution. Layers are connected feed-forwardly, receiving concatenated feature maps from all previous layers to enhance feature reuse and reduce parameters.

In contrast to DenseNet having 3×3 convolutional layers, we increased the number of channels and the kernel sizes used in each dense layer to capture a broader context of the base position, up to 1024 channels and a kernel size of 99 in the last dense block. Each dense block, except the last one, was followed by a 1×1 convolutional layer to reduce features and accelerate processing. The final output was connected to a fully connected layer with a log softmax activation for classification. Our network used connectionist temporal classification (CTC) loss^56^ for gradient descent. Basecalling was performed with a beam search size of 5.

#### Data process and basecaller training

For model training and evaluation, we utilized the Taiyaki (ONT, v5.3.0) to preprocess the training and testing datasets. Each chunk, representing a segment of the raw signal data, contained 4096 signal values. These chunks were normalized using the median absolute deviation method to ensure data consistency. The training set includes data from *Arabidopsis thaliana*, *Homo Sapiens*, *Caenorhabditis elegans*, and *Escherichia coli* from the RODAN^54^ study, using one million signal chunks for training and 100,000 for validation. In addition, we employed a memory optimization technique^57^ to reduce memory consumption during model training.

The test dataset includes datasets from the RODAN study, including *Homo sapiens*, *Arabidopsis thaliana*, *Mus musculus*, *S. cerevisiae* S288C and *Populus trichocarpa*, the published SARS-CoV-2 dataset^11^ and the datasets generated in this study (**Supplementary Data 3, 6**), including *Homo Sapiens*, *P. berghei*, Seneca Valley virus (SVV), Porcine Epidemic Diarrhea virus (PEDV) and Porcine Reproductive and Respiratory Syndrome virus (PRRSV), a total of 10 species (**Supplementary Table 4**). We used PyTorch^94^ (v2.0.1), set the batch size to 32 and trained for 30 epochs. The label smoothing technique of the model and basecalling function were adapted from RODAN^54^.

#### Species-specific model training

The *Mus musculus* dataset from RODAN^54^ was used to train a species-specific model. The training set consists of 20,000 electrical signals, while the validation and testing sets each contained 4,000 electrical signals. The HDF5 data were generated using Taiyaki (ONT, v5.3.0), and species-specific models were trained similarly to the general models, except that the data sources differed. The parameters for training were set to a learning rate of 0.002, a batch size of 32, and the process was conducted over 30 epochs. The nucleotide sequences of the test set were ultimately produced by the DEMINERS’s basecaller utilizing the trained model to ensure accurate representation of the mouse transcripts.

#### Performance Evaluation

Basecallers were evaluated using sequence identity defined as *Accuracy = M / (M+X+I+D)*, where M is the number of matching bases, X is the number of mismatches, I is the number of insertions, and D is the number of deletions. Sequence alignment and accuracy assessment, including quantification of mismatches, insertions, and deletions against a reference genome, were conducted using minimap2^88^ (v2.15, -cs) and the *accuracy.py* function of RODAN^54^ (v1.0).

### RNA extraction from biological samples

#### S. cerevisiae

*S. cerevisiae* W303 was cultured at 30°C in YPD (Yeast Peptone Dextrose) medium containing 20 g/L of glucose, 10 g/L of yeast extraction, and 20 g/L of peptone. The collected cells were centrifuged at 5000 rpm for 5 mins, followed by washing with PBS. The resuspended cells in PBS were frozen with liquid nitrogen and then thawed in 37°C water four times. Total RNA was extracted with the TRIzol reagent (Invitrogen, 15596026) and the mRNA was enriched according to the Dynabeads mRNA purification Kit (Thermo Fisher, 61006).

#### Viruses

Seneca Valley virus (SVV), Porcine Epidemic Diarrhea virus (PEDV), Getah virus (GETV), and Porcine Reproductive and Respiratory Syndrome virus (PRRSV) were provided by the Key Laboratory of Animal Diseases and Human Health of Sichuan Province. The total RNA of these viruses was extracted using TRIzol reagent (Invitrogen, 15596026). Isolation of SARS-CoV-2 RNA was described in the section of ‘Clinical specimen collection and RNA preparation’.

#### Bacteria

*Escherichia coli O157:H7* (ATCC 43895) and *Salmonella enteritid* (ATCC 13076) were cultured in LB media at 37°C to an OD600 of 0.6. Bacteria were collected by centrifugation at 3000 rpm for 5 min and washed with PBS. The resuspended cells in PBS were frozen with liquid nitrogen and then thawed in 37°C water for four times. Total RNA was isolated using TRIzol reagent (Invitrogen, 15596026) following the instructions of the manufacturer, and treated by DNase I (Thermo Fisher, EN0521) at 37°C for 30 min. Poly(A) tailing was performed using *E. coli* Poly(A) Polymerase (NEB, M0276S), and the resulting product was purified using RNA Clean & Concentrator-5 kit (Zymo, R1015) and enriched by the Dynabeads mRNA purification Kit (Thermo Fisher, 61006).

#### Plasmodium berghei

C57BL/6J mice aged between 6–8 weeks were purchased from GemPharmatech (Jiangsu, China) and housed under SPF conditions (Specific Pathogen Free) at the Laboratory Animal Center of West China Second University Hospital. The animal experiments were carried out following the protocols approved by the Institutional Animal Care and Use Committee of West China Second University Hospital [(2018) Animal Ethics Approval No. 024].

Mice were intraperitoneally injected with 10^4^ red blood cells (RBCs) infected with *P. berghei* ANKA parasites. Six days after infection, the blood containing trophozoite stage parasites was collected via cardiac puncture and filtered through leukocyte filters (Bengbu Zhixing Biotech, China). The collected infected RBCs were cultured in IMDM (Thermo Fisher, 12440053) supplemented with 20% fetal bovine serum (Thermo Fisher, 10100147) with 0.5% Penicillin-Streptomycin (Thermo Fisher, 15140122) at 37°C in an atmosphere of 5% CO2 and 5% O2 for 14-16 hours. The maturation of the parasites was examined via Giemsa-stained blood smears. Schizont or late trophozoite stage parasites were separated using nycodenz (Alere Technologies, 1002424) density centrifugation^95^. The upper layer consisted of schizont-infected RBCs and the layer below nycodenz contained a mixture of trophozoite-infected RBCs and uninfected RBCs. The RBCs were lysed with Red Blood Cell Lysis Buffer (Solarbio, R1010), and the parasites were enriched by centrifugation and washed with PBS. Total RNAs of schizont and late trophozoites were extracted using TRIzol reagent (Invitrogen, 15596026) following the manufacturers’ recommendations.

### Clinical specimen collection and RNA preparation

#### Ethics approval

Tumour sample collection and the study design were approved by the Biomedical Research Ethics Committee of West China Hospital (Approval number: 2020.837). The research on COVID-19 specimens was approved by the Biomedical Research Ethics Committee of West China Hospital (Approval number 2020.100, 2020.193 and 2020.267). The swabs were obtained for routine diagnostic purposes, and the remaining RNA samples were provided for research purposes. Written consents were obtained from all patients.

#### Clinical sample collection and RNA extraction

Fresh tumor specimens were collected into OCT from glioma patients undergoing surgical resection at West China Hospital between Sep 2020 to Mar 2021. The total RNA was extracted using the TRIzol reagent (Invitrogen, 15596026). Enrichment of poly(A)+ RNA was performed using Dynabeads™ mRNA Purification Kit (Invitrogen, 61006).

The nasopharyngeal and oropharyngeal swabs were collected from suspected or confirmed SARS-CoV-2 infected individuals, the RNA was extracted from swabs utilizing the RNA extraction kit (Sichuan Maccura Biotechnology, GN7101913). Real-time RT-PCR was performed by amplifying SARS-CoV-2 two target genes, open reading frame 1ab (ORF1ab) and nucleocapsid protein (N), using the 2019-nCoV Nucleic Acid Detection Kit (Sansure Biotech Inc.)

### Nanopore direct RNA sequencing and library preparation

Single-sample direct RNA sequencing (DRS) libraries were prepared using the Direct RNA Sequencing Kit (SQK-RNA002, ONT) following the manufacturer’s protocol (DRCE_9079_v2_revI_14Aug2019, ONT). For multiplexed RNA samples, the DRS libraries were prepared using the DEMINERS approach, which enables the concurrent sequencing of multiple samples within the same run. Sequencing was performed on a PromethION sequencer (ONT) using the R9.4.1 flow cell. Electrical signal data were collected using the MinKNOW software (v21.06.0, ONT).

### Read basecalling, alignment, and visualization

For basecalling of the raw electrical signals contained in FAST5 files, we utilized the DEMINERS basecaller, applying the rna_r9.4.1_hac@v1.0 model, which enabled us to acquire RNA sequence data. The reference genome FASTA files were downloaded from NCBI (https://www.ncbi.nlm.nih.gov). The *P. berghei* genome FASTA and GFF files were downloaded from the PlasmoDB (https://plasmodb.org/plasmo/app). The filtered sequence reads were aligned against the reference genomes (GenBank Accession): *Homo sapiens* (hg38), PEDV (NC_003436), SVV (DQ641257), PRRSV (NC_001961), SARS-CoV-2 (MN908947.3), *E. coil* (NC_000913.3), *S. enterica* (NC_003197.2), *S. cerevisiae* (NC_001133-48), *Prevotella* (NZ_CP019300.1), *Veillonella* (NZ_CABKSO010000001.1), *Streptococcus* (NZ_GL732439.1) and *P. berghei* (PlasmoDB-53, Plasmodb.org), using minimap2^88^ (v2.15-ax map-ont). The direct RNA sequencing reads distribution was assessed by the coverage function of R package GenomicAlignments^96^ (v1.22.1), and visualized by R package ggplot2 (v3.3.6, https://ggplot2.tidyverse.org/). The mutations, deletions and genome coverage were visualized using IGV^97^ (v2.12.2).

### Next-generation sequencing for SARS-CoV-2 samples and data process

The RNA extracted from nasopharyngeal and oropharyngeal swabs was subjected to a PCR enrichment of the SARS-CoV-2 genome using the SARS-CoV-2 Full Length Genome Panel (Genskey, 2205) following the manufacturer’s instructions. The cDNA was then used to prepare paired-end sequencing libraries through a ligation method (Genskey, 2205). According to the manufacturer’s recommendations, the resulting libraries were sequenced on an MGISEQ-2000 platform using the sequencing reaction kit (Genskey, GS-2000-FCS-SE100).

The low-quality regions, adaptor sequences, and sequencing primers were trimmed using fastp^98^ (v0.23.2). The clean reads were mapped to the reference genome (MN908947.3) by Bowtie2^99^ (v2.4.4). After alignment, all mapped reads were aggregated to construct a consensus sequence employing methods from prior research^100^.

### Identification of SNVs and deletions

For DEMINERS, we selected alignment reads from SVV, PRRSV, SARS-CoV-2, and glioma samples. For NGS, we selected alignment reads from SARS-CoV-2, and for glioma samples, we used previously published cDNA-NGS data^101^ (Genome Sequence Archive, accession ID HRA001865). These reads were aligned to the reference genome (hg38) using Bowtie2^99^ (v2.4.4). Subsequently, we generated VCF files containing mutation information using the mpileup function in Samtools^102^ (v1.9) and the call function in bcftools^103^ (v1.8). Variants with a quality score below 20 were excluded. Deletions were identified based on a minimum expression level of 3, deletion length greater than 3 bp, and a deletion coverage greater than 30%.

### Isoform reconstruction and quantification

#### Read alignment and correction

The reads were aligned to the reference genome using minimap2^88^ (v2.15, -ax splice -k14 -uf –secondary= no). Annotated splice junctions were provided to guide the alignment. The alignment output was subsequently sorted and converted into the BAM format using Samtools^102^ (v1.9). Then, TranscriptClean^104^ (v2.0.2) was used to correct mismatches, insertions, deletions, and non-canonical splice junctions, using known variants and splice junctions as references. The cleaned reads were next realigned to the reference genome for subsequent analysis.

#### Isoform identification and quantification

The process of isoform identification was performed using a custom-developed R script (https://github.com/LuChenLab/DEMINERS/tree/main/scripts). This process began with reading BAM files and filtering reads based on the mapping quality with a minimum MAPQ score using Rsamtools (v2.6.0, https://bioconductor.org/packages/Rsamtools). Subsequently, it identifies novel splice junctions supported by user-defined read count thresholds using GenomicAlignments^96^ (v1.22.1). Strand-specific analysis is also incorporated to delineate transcription directionality. Several parameters were configured, such as the minimum length of insertions or deletions and the minimum reads for various novel transcription features. Moreover, multi-core processing was enabled by using doParallel (v1.0.16, https://CRAN.R-project.org/package=doParalle). This workflow generated a FASTA file with reconstructed isoform sequences, a JSON file mapping reads to corresponding transcripts, and an annotated GTF file cataloging the novel isoforms.

Quantification was achieved via an Expectation-Maximization (EM) algorithm after isoform identification. Initially, the EM algorithm assigned a compatibility score to each isoform based on the proportion of isoform per read. During the EM cycles, isoform abundances were calculated by aggregating compatibility scores for each read aligned to the isoform and normalizing by the total compatibility scores for all alignments. The compatibility index for each read-isoform pair was updated by dividing the isoform abundance by the sum of abundances for reads aligned to that isoform. The EM iterations continue until the predefined convergence criterion is met, indicating minimal changes in cumulative isoform abundance across successive EM rounds. Upon convergence, the algorithm yields an accurate isoform abundance table from which estimated counts were derived, providing a precise measurement of isoform presence across samples based on the demultiplexing method.

### Splice events identification

The splice junction (SJ) identification and quantification were performed using the junctions function from R package GenomicAlignments^96^ (v1.22.1) The alternative splicing events were extracted from aligned bam files using SUPPA2^105^ (v2.3).

### Prediction and analysis of protein structures from isoform sequences

Possible open reading frames (ORFs) were predicted based on the isoform nucleotide sequences using R package ORFhunteR^106^ (v1.4.0). The longest ORF was translated into an amino acid sequence and subjected to structural prediction using AlphaFold^107^ (v3). Functional predictions were performed with InterProScan5^108^ (v5.68-100.0). Finally, protein structure alignment and visualization were conducted using PyMOL (v3.0, https://www.pymol.org/).

### Assessing prognostic value of genes and splice junctions in gliomas

To verify whether *GFAP* splicing events are regulated by *METTL3*, we downloaded the RNA-seq on K562 cells treated with a CRISPR gRNA against *METTL3* from the ENCODE database^74^ and identified the expression of *GFAP*-associated SJs. To explore the prognostic value of genes and splice junctions in gliomas, we downloaded the gene expression and clinical data of glioblastoma multiforme (GBM) from the TCGA Data Portal (https://tcga-data.nci.nih.gov/tcga/), and linear junctions expression data from the RJunBase database^75^. Using the R packages survminer (v0.4.9, https://CRAN.R-project.org/package=survminer) and survival (v3.2.13, https://CRAN.R-project.org/package=survival), we determined the optimal cutpoints for continuous variables, plotted survival curves, and computed p-values to compare survival curves.

### m^6^A identification and quantification

Raw current signal data was processed by re-squiggled algorithm with Tombo (Oxford Nanopore Technologies, v1.5.1) for accurate reference alignment. Nanom6A^13^ (v2021.10.22) extracted statistical metrics from Nanopore signals for m^6^A modification probability post-re-squiggle. At the same time, read IDs can be used to associate modification sites from Nanom6A^13^ (v2021.10.22) with isoform. In parallel, TandemMod^63^ (v2023.08.18) performed feature extraction and m^6^A modified ratio calculations based on designated motifs. Simultaneously, m6Anet^62^ (v2.1.0), with signal segmentation by Nanopolish^4^ (v0.13.2), predicted m^6^A modifications, employing event scaling and signal indexing for feature extraction.

### Association between SNV and m^6^A Site in SARS-CoV-2 genomes

Firstly, we identified m^6^A sites in two samples with over 50 SARS-CoV-2 reads using three software tools: Nanom6A^13^ (v2022.12.22), TandemMod^63^ (v2023.08.18) and m6Anet^62^ (v2.1.0). A site was deemed highly credible if identified more than five times by all three tools in at least two samples (**Supplementary Data 5**). From these m^6^A sites, we extended a 500 bp window both upstream and downstream. SNV sites, identified in the NGS data of the corresponding samples, were subsequently intersected with these m^6^A sites and their surrounding windows, followed by a calculation of SNV frequencies. Additionally, 300 random genomic sites without any SNVs were selected from the SARS-CoV-2 genome to serve as control random backgrounds.

### Differential analyses of malaria parasites

We utilized DESeq2^109^ (v 1.32.0) to identify differentially expressed genes and isoforms (Wald-test, *P* value < 0.05). Nanopolish^4^ (v0.13.2) was used to estimate the lengths of poly(A) tails. The Wilcoxon test was used to determine the significant changes in poly(A) tail length between the trophozoite and schizont stages of malaria parasites. The differential m^6^A ratios were determined using T-tests with a significance cutoff of *P* value < 0.05.

### Metagene motif distribution analysis

MetaplotR^110^ (v1.0) was used to analyze the motifs of m^6^A peaks in DRS. The motif closest to DRACH was identified based on the annotation information of the reference genome downloaded from the UCSC Genome Browser (http://hgdownload.soe.ucsc.edu/). The relative localization of the motif coordinates in the transcript regions (5’UTR, CDS, 3’UTR) was determined. The relative lengths of the three transcript regions were defined by scaling genes that contain at least one m^6^A peak.

### Metagenomic analysis

The demultiplexed DRS data were filtered according to the criteria of qual > 5 and length > 100. The U bases were converted to T bases for further analysis. The taxonomic classification and calculation of taxonomic abundance were performed using BugSeq^111^ (v2023-11-27) with a default metagenomic database. For domain-level analysis, reads from all samples were pooled to identify and calculate the percentage of each domain. For genus discrimination, reads from nasopharyngeal and oropharyngeal swabs were analyzed separately, with each swab type showing the percentage of the top ten genera. For species discrimination, species with read counts above 10 were retained, and DESeq2^109^ (v1.32.0) was used to identify species that exhibited differential distribution between nasopharyngeal and oropharyngeal samples (Wald-test, P value < 0.05).

### Genome assembly

For the RNA virus samples, including SVV, PRRSV, and metagenomic samples from COVID-19, we performed genome assembly using the following method. First, we utilized Canu (v2.2)^112^ with default parameters to assemble the genomes. To assess the consistency with reference genomes, we used FastANI (v1.1)^113^ to calculate the average nucleotide identity between the assembled genomes and the reference genomes. For the COVID-19 metagenomic samples, we used BLASTn (v2.12.0)^114^ to map the assembled contigs to the NCBI non-redundant nucleotide database (nr/nt) for taxonomic annotation. We set the parameters *num_alignments* and *max_hsps* to 1 to identify the best match. Finally, we used TaxonKit (v0.16.0)^115^ to complete the full taxonomic classification based on the respective TaxIDs.

### Sanger sequencing validation

For PRRSV samples, primers spanning the genome region from 13941 to 15340 were designed to verify all point mutations within that region. The primer sequences were listed in **Supplementary Data 3.** For glioblastoma samples, primers spanning four-point mutations were designed. The primer sequences were listed in **Supplementary Data 6.** Reverse transcription (RT) was performed using SuperScript IV (ThermoFisher Scientific, 18090050) with random hexamers. Amplification was performed using the KAPA HiFi HotStart PCR Kit (Roche, KK2501), and the sequences were determined by Sanger sequencing.

## Data and Code Availability

The raw and processed sequencing data generated in this study were submitted to the NCBI BioProject database (Accession number PRJNA911167).

The software and related scripts of DEMINERS were uploaded to GitHub (https://github.com/LuChenLab/DEMINERS).

The pre-trained models for basecalling models were deposited in Figshare (https://figshare.com/articles/dataset/Densecall_models/25712856).

The pre-trained models for barcode demultiplexing were deposited in Figshare (https://figshare.com/articles/online_resource/DecodeR_Models/22678729).

## Author contributions

LC, J-wL and JG conceived the project and designed the experiments. JS, CC developed a multiplexing experimental protocol and conducted the experiments. CT developed the demultiplexing pipelines. LL developed the basecalling method. YW, BY and BC assisted by JZ, JJ and JW provided the clinical samples. JS and CC assisted by YZ, H-cW, GS and MC prepared the RNA and conducted the direct RNA sequencing. JS and CT assisted by QY, DZ, KL, ZX, TC, ZH, DL, WZ and LC performed the bioinformatic analysis. LC, J-wL and JG analyzed the results. JS, LC and J-wL prepared the manuscript. JS, CT, and CC assisted by QY, DZ, ZX, YZ, and WZ generated the figures and helped prepare the manuscript.

## Acknowledgments

This work was supported by the National Natural Science Foundation of China (82341122, 92369116, 82370233, 82300133, 81722004, and 31800772), Sichuan Science and Technology Program (2021YFS0027), National Key Research and Development Program of China (2017YFA0106800 and 2021YFF0702000), and China Postdoctoral Science Foundation (2023M742455). We thank Dr. Xinqiong Li for providing us with the virus samples. We thank Coseque (Chengdu, China) for providing support in basecalling algorithm.

## Supplementary Tables

**Supplementary Table 1.** Comparative performance of demultiplexing direct RNA-seq data using different machine-learning algorithms.

**Supplementary Table 2.** Accuracy and performance of DEMINERS in the training and test sets.

**Supplementary Table 3.** Comparative analysis of DEMINERS and established demultiplexing methods.

**Supplementary Table 4.** Median performance of different basecallers in basecalling direct RNA-seq data.

## Supplementary Data

**Supplementary Data 1.** Data used to establish DEMINERS barcode classification algorithm.

1. Sequences and usage of RNA transcription adapters and barcodes.
2. Sequences of in-vitro transcribed RNA and the related primer sequences.
3. Information of five sequencing runs.
4. Data used for training or testing for different machine-learning algorithms.
5. Barcodes and reads used in demultiplexing test of DEMINERS.
6. Data used for model training or testing of DEMINERS in classifying different numbers of barcodes
7. Accuracy and recovery rates of DEMINERS on the test set at different probability cutoffs.

**Supplementary Data 2.** Data used to evaluate DEMINERS performance.

1. Data used to demultiplex four DeePlexiCon or four Poreplex barcodes.
2. Accuracies of DEMINERS in demultiplexing four DeePlexiCon barcodes.
3. Data used to demultiplex 24 custom-designed barcodes.
4. Accuracies of DEMINERS and DeePlexiCon in demultiplexing 24 barcodes.

**Supplementary Data 3.** DEMINERS performance on pathogen identification.

1. Sample information and DEMINERS retrieved reads.
2. Accuracies of DEMINERS on pathogen identification.
3. Primers used for Sanger sequencing.
4. Identification and validation of SNVs in PRRSV genome.

**Supplementary Data 4.** DEMINERS performance on clinical metagenomics.

1. Information of 24 swab samples for COVID-19 testing.
2. Taxonomic assignment and relative abundance.
3. Metagenomic contig assembly and taxonomic classification.
4. m^6^A sites identified using three software.
5. Identification and validation of SNVs and deletions.

**Supplementary Data 5.** DEMINERS applied on parallel comparisons of RNA features.

1. Information of *P. berghei* samples and retrieved reads.
2. Analysis of splice junctions, isoforms, and m^6^A of different stages of *P. berghei*.
3. Prediction of coding sequence and protein sequence of novel isoforms.

**Supplementary Data 6.** DEMINERS performance on glioma samples.

1. Information of glioma samples.
2. Read numbers and mutation sites identified by NGS and DEMINERS.
3. Primers used for Sanger sequencing.

---

**Extended Data Fig 1.**
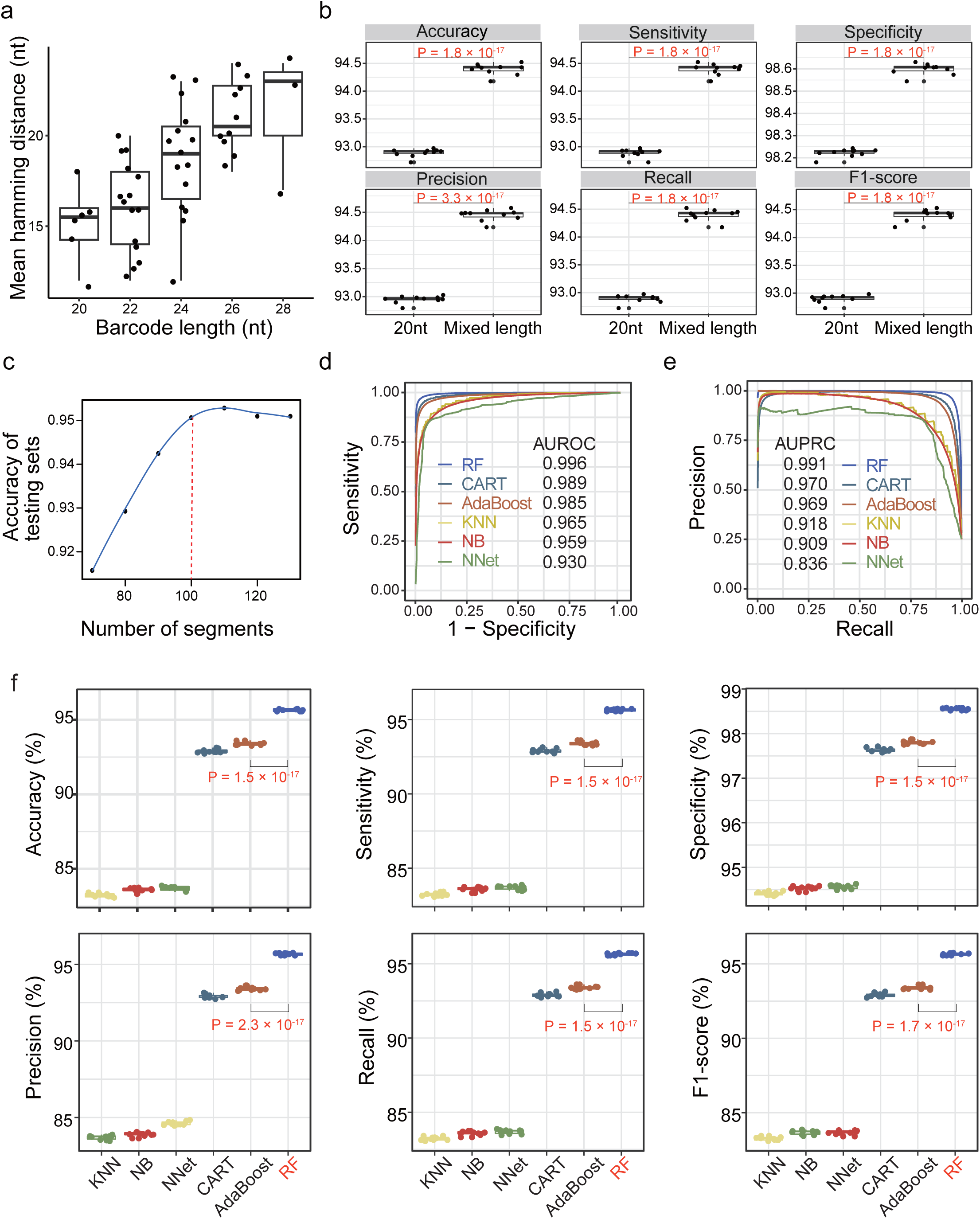
Performance in classifying different barcode configurations and comparisons of 6 machine learning-algorithms. **a,** Box plot illustrating the correlation between average Hamming distance and different barcode lengths. **b,** Box plots comparing the demultiplexing performance of uniformed 20-nt barcodes versus barcodes of mixed length. P values, Wilcoxon test. **c,** Accuracy curve showing the relationship between the number of signal segments used in testing set, with the optimal number (100) indicated by a red dashed line. **d,** Receiver Operating Characteristic (ROC) curves assessing the performance of six different machine0learning methods. AUROC, the area under the ROC curve. **e,** Precision-Recall (PR) curves for the same methods. AUPRC, the area under the PR curve. **f,** Comparisons of demultiplexing performance of 6 machine-learning methods, measured by accuracy, sensitivity, specificity, precision, recall, and F1-score. P values, T-test. RF, random forest (red); CART, classification and regression tree; KNN, k-nearest neighbor; NB, naïve Bayes; NNet, neural network.

**Extended Data Fig 2.**
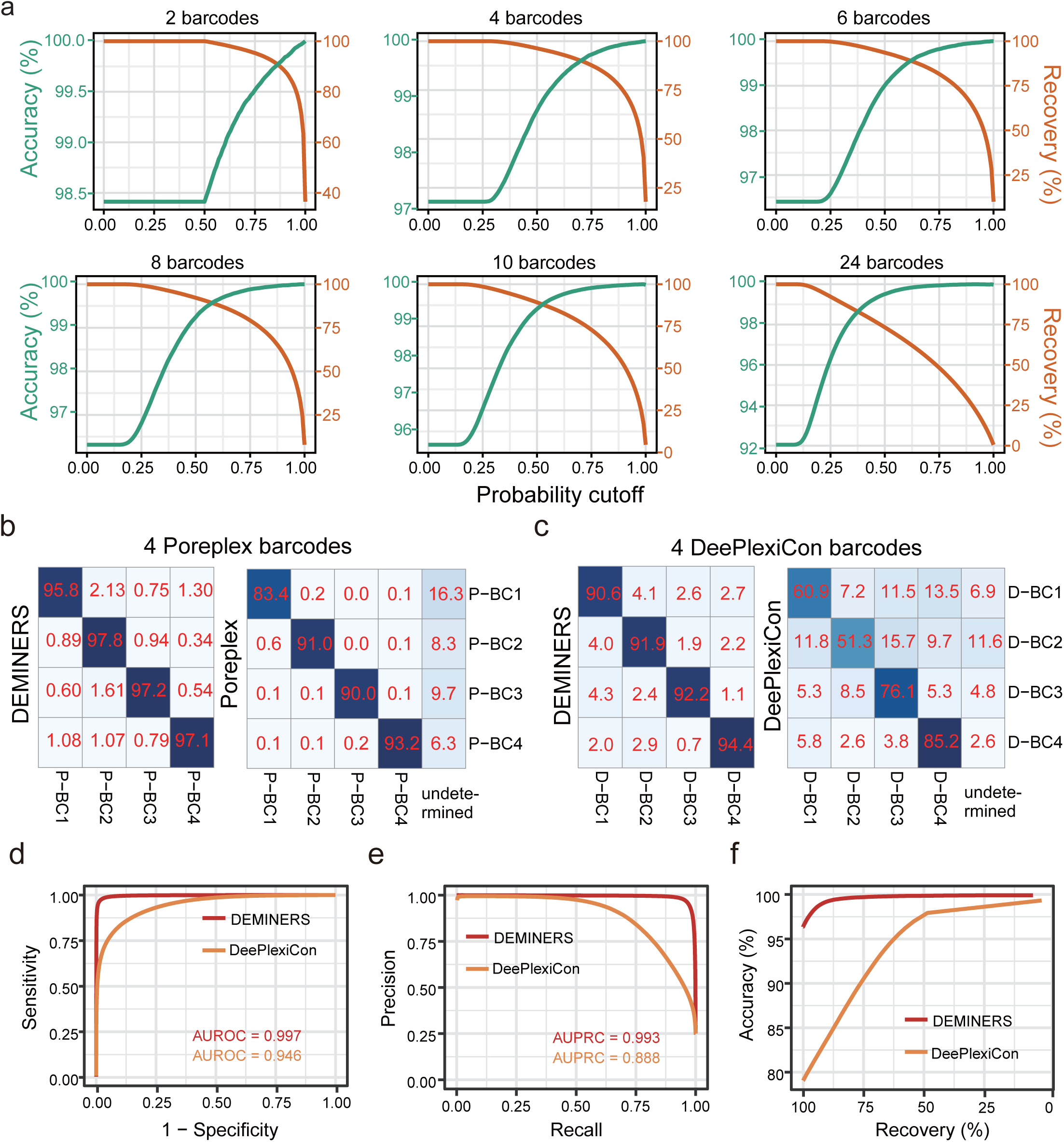
Performance comparison of DEMINERS, Poreplex and DeePlexiCon. **a,** Accuracy and recovery rate curves for different numbers of barcodes showing the impact of probability cutoffs on predictive performance. **b,** Confusion matrices comparing the performance of DEMINERS and Poreplex^57^ classifying 4 Poreplex barcodes. **c,** Confusion matrices comparing DEMINERS and DeePlexiCon^58^ classifying 4 DeePlexiCon barcodes. **d,** ROC curves comparing DEMINERS and DeePlexiCon classifying 4 DeePlexiCon barcodes. **e,** PR curves comparing DEMINERS and DeePlexiCon classifying 4 DeePlexiCon barcodes. **f,** Accuracy-recovery rate trade-offs for DEMINERS and DeePlexiCon classifying 4 DeePlexiCon barcodes.

**Extended Data Fig 3.**
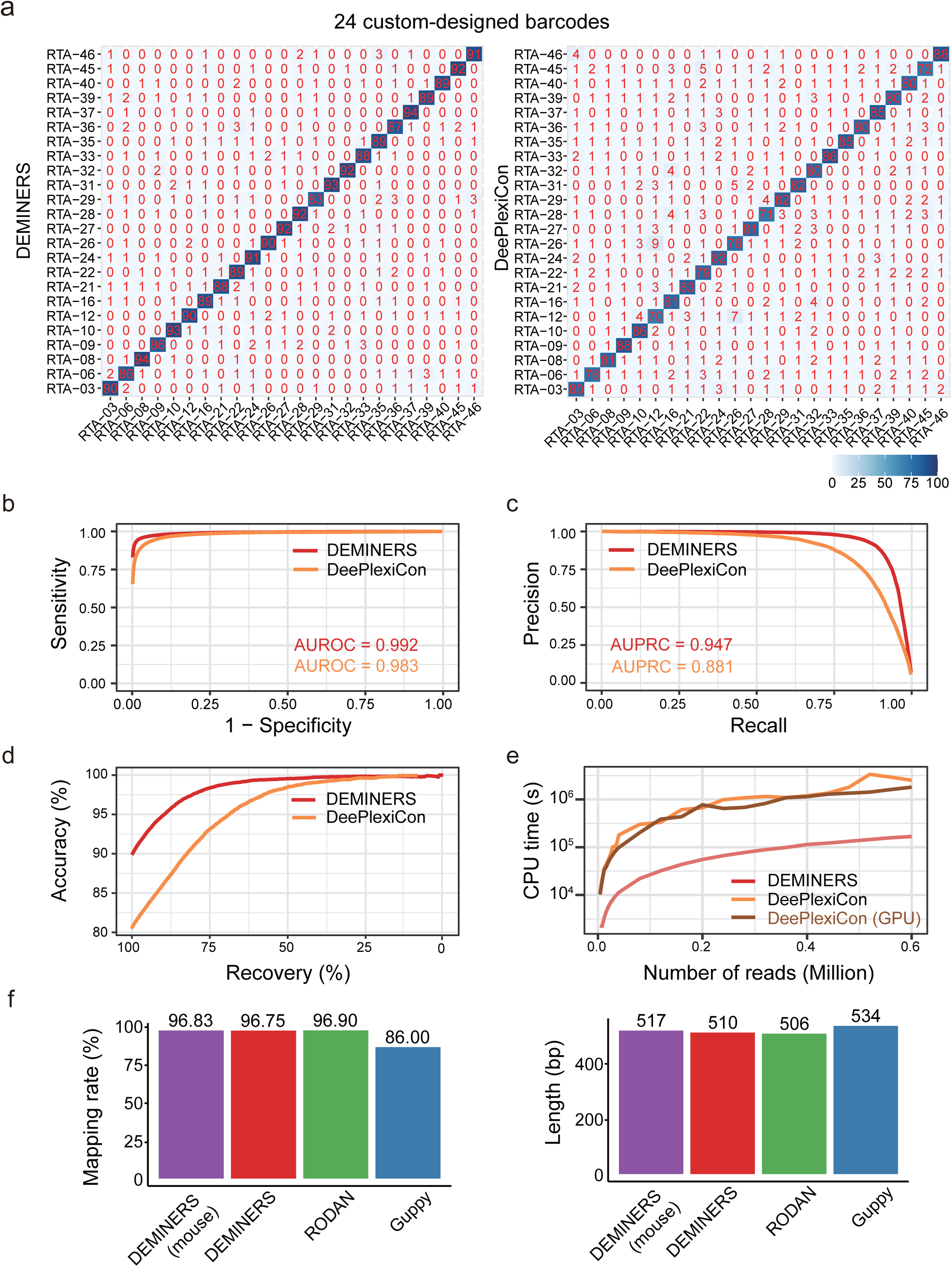
Performance of demultiplexing and basecalling by different methods. **a,** Confusion matrices comparing demultiplexing performance of DEMINERS and DeePlexiCon^58^ using 24 custom-designed barcodes. **b-d,** Comparative analysis of DEMINERS and DeePlexiCon classifying 24 custom-designed barcodes. ROC (**b**) and PR curves (**c**) and accuracy-recovery rate trade-offs (**d**) are shown. **e,** CPU running time for increasing numbers of reads for DEMINERS, DeePlexiCon, and the GPU of DeePlexiCon. **f,** Performance metrics of accuracy, mapping rate and read length for DEMINERS (mouse-specific and general modes), RODAN, and Guppy.

**Extended Data Fig 4.**
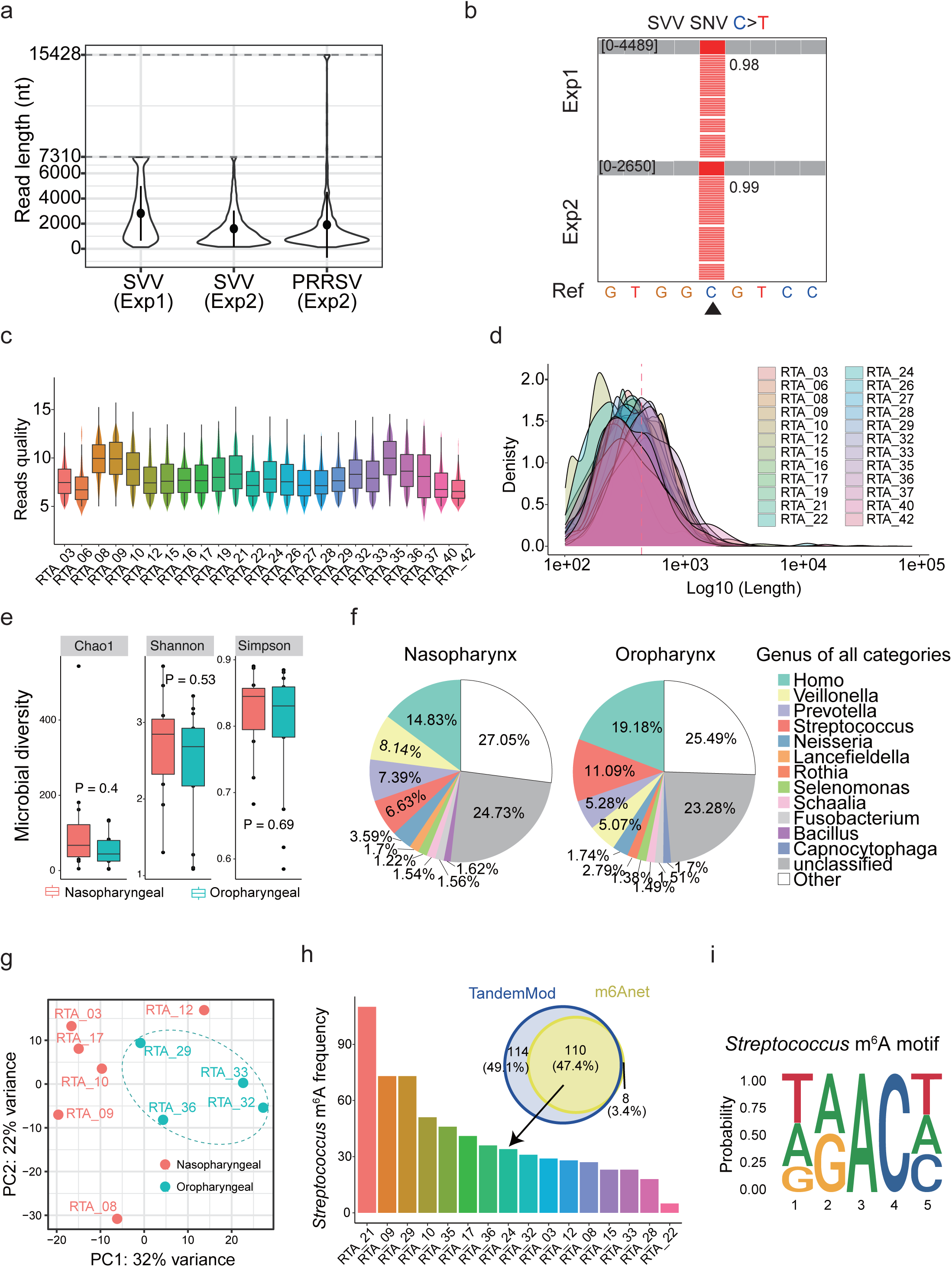
Performance of DEMINERS in pathogen RNA analysis. **a,** Distribution of read lengths in SVV and PRRSV samples, with reference genome lengths indicated by dashed lines. **b,** IGV visualization of a SNV (C-to-T at position 5700) in SVV genome identified by DEMINERS in two experiments, with the SNV indicated by a triangle. The numbers represent the read frequency of the SNV. **c,** Box plot depicting the read quality across 24 swab samples. **d,** Density plot reflecting lengths of 24 swab samples, log10-transformed on the x-axis, with a mean length indicated by a red dashed line. **e,** Box plots showing microbial diversity (Chao1, Shannon, and Simpson indices) in nasopharyngeal and oropharyngeal swabs. P values, Wilcoxon test. **f,** Pie charts depicting the distribution of genus identified in nasopharyngeal and oropharyngeal swabs. **g,** PCA plot visualizing sample grouped by microbial read numbers. **h,** Venn diagram representing m^6^A sites identified by TandemMod^71^and m6Anet^70^ for *Streptococcus*, and the bar graph showing m^6^A frequency in 16 swab samples. **i,** Sequence logo of identified m^6^A site motifs for *Streptococcus* in 16 swab samples.

**Extended Data Fig 5.**
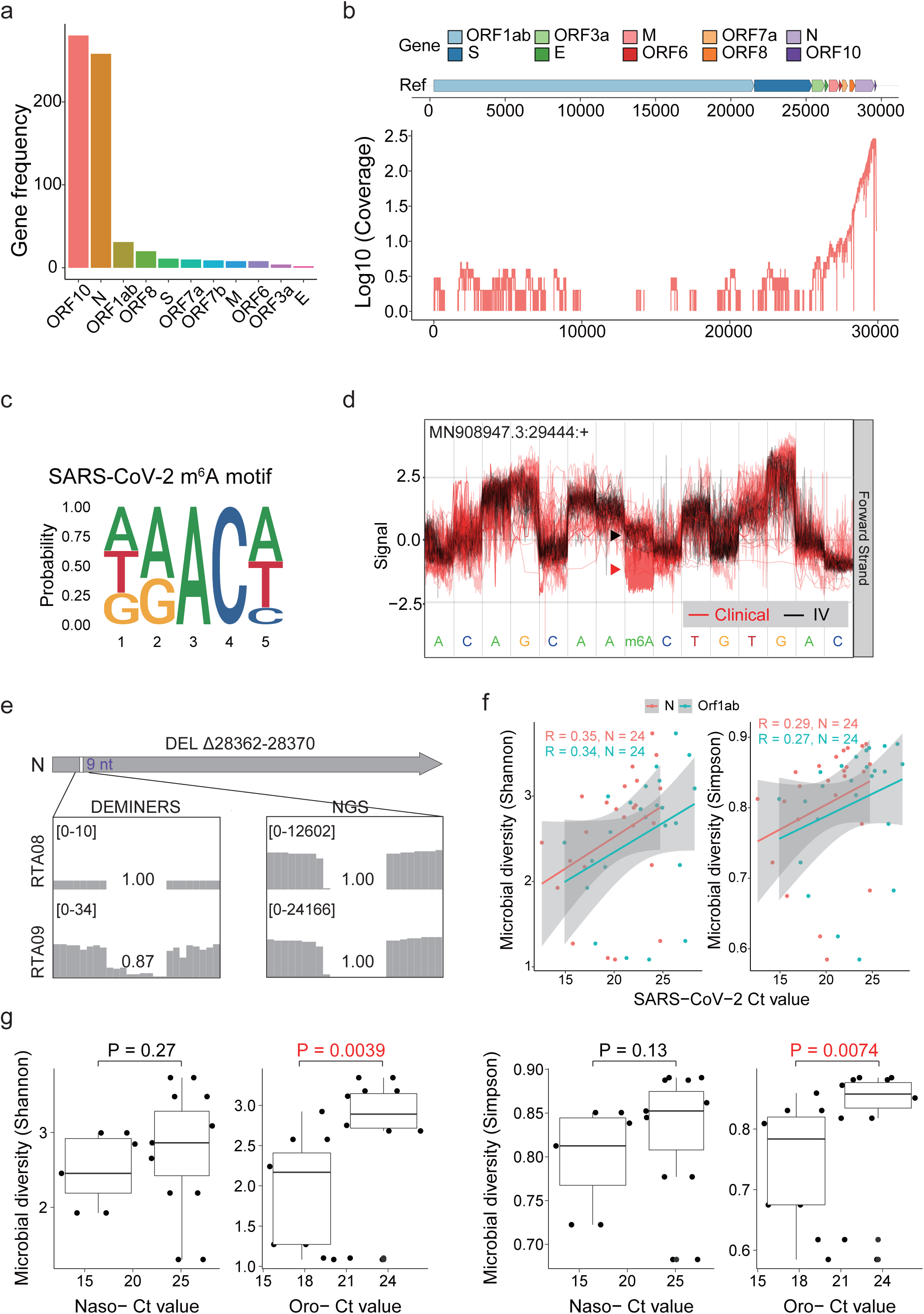
SARS-CoV-2 RNA and microbial diversity analyzed by DEMINERS. **a,** Bar plot showing the frequency of SARS-CoV-2 genes identified in all detected samples (n=14). **b,** Density plot illustrating the genome coverage of SARS-CoV-2 reads in all detected samples (n=14), log10-transformed on the y-axis. **c,** Sequence logo of identified m^6^A site motifs for SARS-CoV-2 in all detected samples (n=14). **d,** Distinct ionic current signals indicating RNA modifications at position 29,444. Red lines, SARS-CoV-2 from swab specimens (clinical); black lines, SARS-CoV-2 maintained in vitro culture (IV). **e,** IGV visualization of a 9-bp deletion (28362 to 28370) identified by DEMINERS and NGS in two samples. The numbers represent the average deletion frequencies. **f,** Pearson correlation between Ct value of SARS-CoV-2 *N* and *Orf1ab* genes and microbial diversity (Shannon or Simpson). R, Pearson correlation coefficients; N, sample size. **g,** Box plots showing the microbial diversity in nasopharyngeal (Naso-) or oropharyngeal (Oro-) swabs with high-Ct value (Ct>21) or low-Ct value (Ct≤21). Microbial diversity was determined by Shannon (left panel) or Simpson (right panel). P values, Wilcoxon test.

**Extended Data Fig 6.**
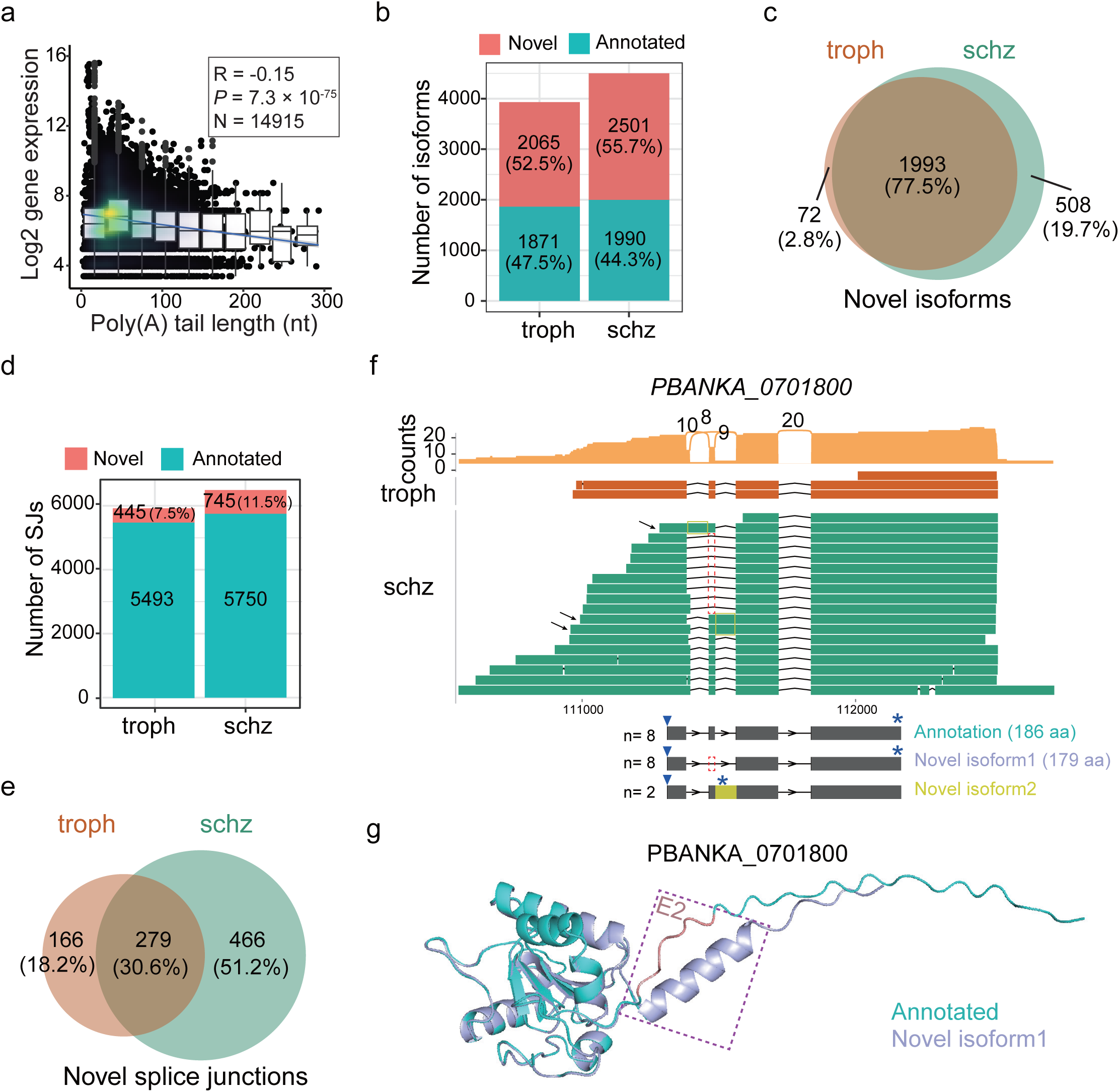
Post-transcriptional mRNA processing in malaria parasites. **a,** Scatter plot showing Pearson correlation between log2 gene expression and poly(A) tail lengths in all malaria samples. R, Pearson correlation coefficient; P, relative P value; N, relative sample size. **b,** Bar plots showing the numbers and percentages of novel and annotated isoforms in trophozoites and schizonts. **c,** Venn diagram comparing novel isoforms in trophozoites and schizonts. **d,** Bar plots showing the number of novel and annotated splice junctions in trophozoites and schizonts. **e,** Venn diagram comparing novel splice junctions between trophozoites and schizonts. **f,** Sashimi plot and read alignments for *PBANKA_0701800* in trophozoites and schizonts. The red dashed box indicates the skipped exons, the yellow box indicates the retained introns. The arrows in the left indicate the reads of novel isoforms. Annotated transcript and novel isoforms with more than 2 DRS reads are shown below. The inversed triangles indicate the predicted start sites the stars indicate the predicted stop site and the numbers in the brackets showing the length of the predicted translated proteins. **g,** Pymol visualization of predicted protein structures based on the annotated and novel isoform. The purple dashed box highlights structural differences.

**Extended Data Fig 7.**
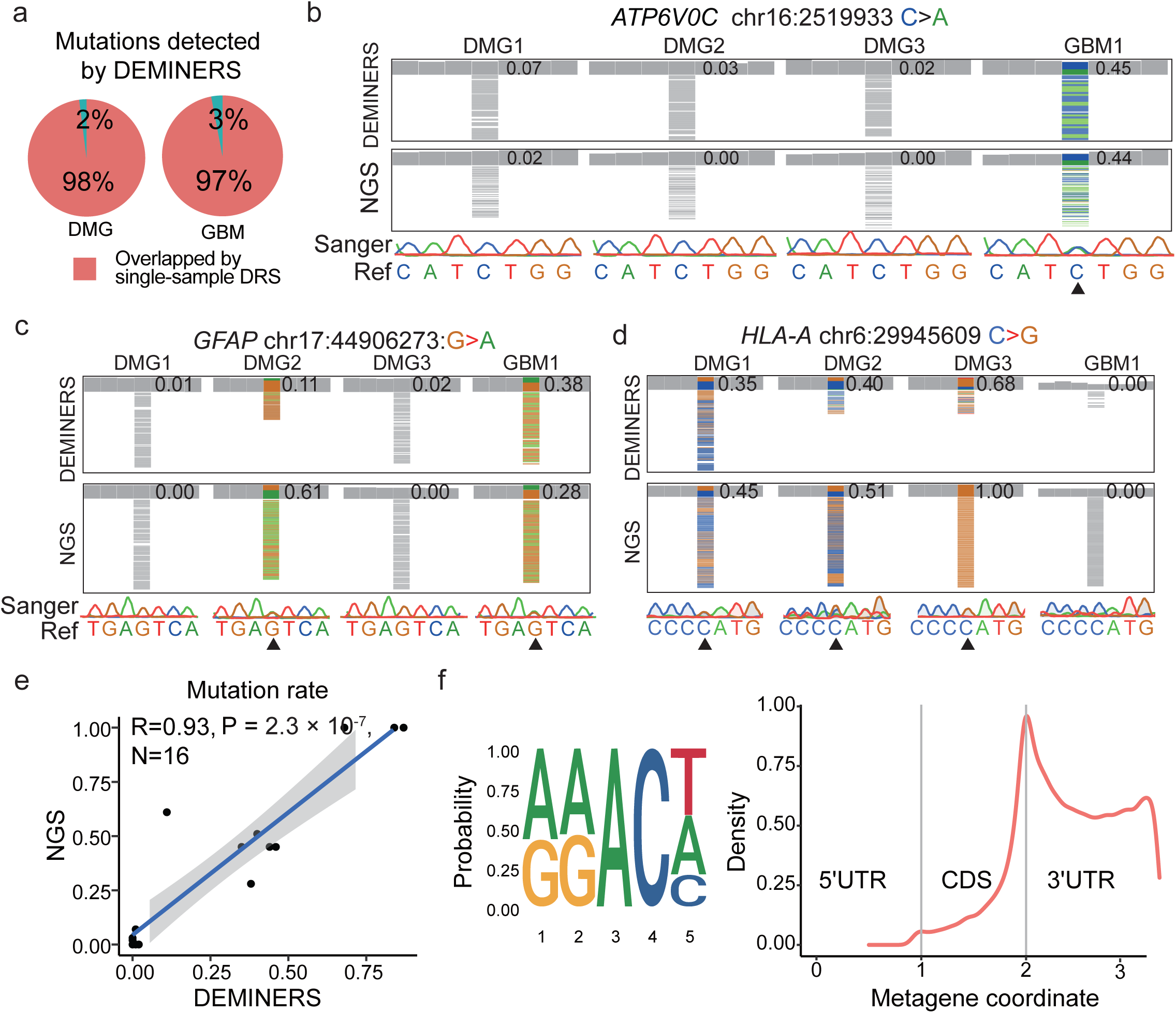
Mutations and m^6^A analysis of human glioma samples. **a,** Pie charts showing the percentage of mutations identified by DEMINERS and single-sample DRS in DMG and GBM samples. **b-d,** IGV visualization of three mutations: C-to-A at chr16 2519933 in *ATP6V0C* **(b)**, G-to-A at chr17 44906273 in *GFAP* **(c)**, C-to-G at chr6 29945609 in *HLA-A* **(d)**. The numbers represent the read frequencies of the mutations. The reference sequence (Ref) and Sanger sequencing chromatograms. **e,** Scatter plot showing Pearson correlation of the 4 mutation rates between DEMINERS and NGS verified by Sanger sequencing in 4 samples. R, Pearson correlation coefficient; P, relative P value; N, relative sample size. **f,** Sequence logo of identified m^6^A motifs (**left**) and distribution of m^6^A sites across transcriptomic regions (**right**) in 4 glioma samples.

